# IL-31 uncouples skin inflammation from itch sensation in allergic dermatitis

**DOI:** 10.1101/2021.05.12.443916

**Authors:** Marlys S. Fassett, Joao M. Braz, Carlos A. Castellanos, Andrew W. Schroeder, Mahsa Sadeghi, Darryl J. Mar, Connie J. Zhou, Jeoung-Sook Shin, Allan I. Basbaum, K. Mark Ansel

## Abstract

Despite a robust literature associating IL-31 with pruritic inflammatory skin diseases, its influence on cutaneous inflammation and on the interplay between inflammatory and neurosensory pathways remain unmapped. Here, we examined the effects of IL-31 and its receptor IL31RA on both inflammation and pruritus in mouse models of dermatitis, including chronic topical house dust mite (HDM) exposure. Unexpectedly, *Il31* deficiency increased cutaneous adaptive type 2 cytokine-producing cells and serum IgE. In addition, M2-like macrophages capable of fueling feedforward pro-inflammatory loops were selectively enriched in *Il31ra*-deficient skin. Thus, IL-31 is not strictly a pro-inflammatory cytokine, but rather an immunoregulatory factor that limits the magnitude of allergic skin inflammation. In contrast, *Il31*-deficient mice displayed a deficit in HDM-induced scratching. Itch reduction occurred despite intact – and in some cases increased – responsiveness of sensory neurons to other pruritogens released during HDM challenge, highlighting the non-redundant contribution of IL-31-receptive sensory afferents to pruritus in environmental allergen-induced dermatitis. When present, therefore, IL-31 uncouples circuits driven by sensory neurons and immune cells that converge in inflamed skin.

## INTRODUCTION

Many cytokines are produced during type 2 skin inflammation. In this milieu, the IL-6/gp130-family cytokine IL-31 stands out for its ability to directly engage its heterodimeric receptor, IL31RA/OSMRβ, on both myeloid cells and on cutaneous itch-provoking sensory neurons (pruritoceptors)(*1, 2*). That dual expression pattern links skin inflammation with neurosensory pathways that mediate itch, and elevates IL-31 as a promising therapeutic target in a growing number of itchy inflammatory skin diseases including atopic dermatitis (AD)(*3*) and prurigo nodularis(*4*).

Transgenic mice that constitutively overexpress *Il31* in T cells (IL31Tg) develop spontaneous itch so severe that they develop scratching-induced skin lesions(*1*), demonstrating that T cell-derived IL-31 is sufficient to trigger pruritus. Scratching induced by hapten-mediated dermatitis is ameliorated in *Il31*-deficient animals, confirming that *Il31* is necessary for contact hypersensitivity-associated itch(*5*). As intradermal or intrathecal administration of recombinant IL-31 induces acute-onset scratching(*2, 6*), it is likely that the pruritogenic effect of IL-31 results from direct activation of IL31RA/OSMRβ on a subset of non-histaminergic pruritoceptive afferents marked by the calcium channel TRPV1(*2*).

The skin pathology that develops spontaneously in IL31Tg animals also suggests a pro-inflammatory role for IL-31(*1*). In IL31Tg animals, however, the contributions of IL-31 to tissue inflammation are difficult to discern because effects of IL-31 on cutaneous immune cells and keratinocytes cannot be dissociated from secondary effects of scratching-induced skin injury. In fact, enhanced rather than diminished type 2 inflammatory responses were observed in lung and gut in *Il31ra*-deficient mouse strains, indicating that IL-31 can function as a negative regulator of type 2 inflammation(*7–9*). There are multiple potential explanations for these seemingly-discordant results. First, IL31Tg skin and *Il31ra*-deficient lung and gut phenotypes may reflect physiological differences specific to each organ(*9*); without scratching-induced skin injury as a confounder, *Il31ra*-deficiency may unmask inhibitory effects of IL-31 on tissue inflammation. It has also been suggested that amplified type 2 inflammatory phenotypes observed in *Il31ra*-deficient animals may occur as a consequence of liberating its binding partner, OSMRβ, and thus enhancing signaling via OSMRβ’s other ligands(*10*). A final, untested model could invoke differential contributions of the IL31RA+ myeloid versus IL31RA+ neural pathways engaged by IL-31. For example, in some patients and/or disease states, circuits activated by IL-31-dependent sensory afferents may dominate over those mediated by cutaneous inflammatory cells (e.g. prurigo nodularis), or vice versa.

Whereas IL-31-dependent itch-sensory pathways have been well-characterized (*11, 12*), the contributions of IL-31 to cutaneous inflammation remain unclear. It is particularly important to address this knowledge gap in the context of chronic AD and its variants, the clinical context in which *Il31* expression is most often detected(*13, 14*). Here we report the consequences of perturbing both myeloid and neurosensory IL-31-dependent pathways by inducing chronic allergic skin inflammation in *Il31-*deficient mice.

## RESULTS

### Generation and validation of IL31KO mice

We generated *Il31-*deficient mice from *Il31^tm1e(EUCOMM)Wtsi^* C57Bl6/N ES cells provided by the European Conditional Mouse Mutagenesis (EUCOMM) Program(*15*) (Fig. 1A and S1). Consistent with previous reports(*1, 16*), we detected *Il31* mRNA in *in vitro*-differentiated Th1 and Th2 cell subsets from C57Bl/6 wildtype (WT) mice (Fig. 1B) and used this *in vitro* system to confirm that *Il31* mRNA was undetectable in CD4 T cells from *Il31^null/null^* mice (hereafter ‘IL31KO’), validating the mutant allele. Similarly, intracellular staining with an IL-31-specific antibody detected robust IL-31 production in restimulated WT Th1 and Th2 cultured cells, but none in IL31KO cultures (Fig. 1C). Importantly, expression of lineage-defining cytokines in Th1 (IFNγ) and Th2 (IL-4, IL-13) cells was unaffected by *Il31* deficiency (Fig. 1B-C), indicating that lineage specification of Th cells does not depend on IL-31. In Th2 cultures, backgating analysis on IL-31 positive cells demonstrated that IL-31+ cells were enriched in IL-4/IL-13 production (Fig. 1D), highlighting IL-31 production as part of an effector cytokine program.

**Fig. 1.**
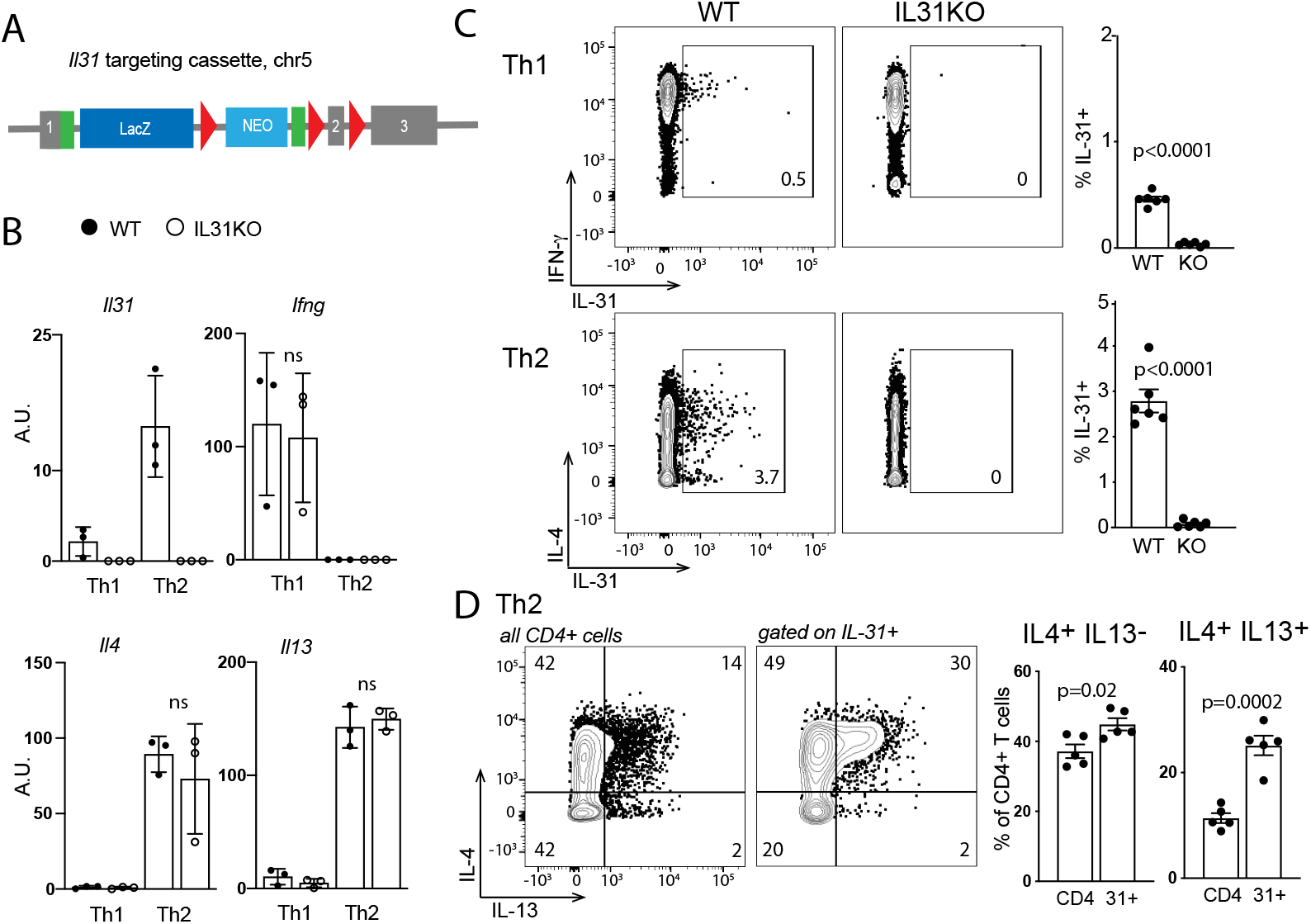
Design and validation of *Il31*^KO^ animals. **(A)** Schematic of the *Il31* targeting construct (*Il31*^tm1e(EUCOMM)Wtsi^, MGI 1923649). The transgene interrupts the endogenous 3-exon *Il31* locus (mm10 ch5:123480419-123482049, minus strand) at position 123481232, introducing loxp sites flanking exon 2. Genetic cross of this transgenic mouse to a β-actin Cre recombinase-expressing strain results in a frameshift mutation and generation of a nonsense transcript (see also Fig. S1). **(B)** Expression of *Il31, Ifng, Il4,* and *Il13* by Taqman qPCR. RNA was extracted from *in vitro*-differentiated Th1 and Th2 WT and IL31KO lymph node CD4 T cells. Black circles (WT), open circles (IL31KO). **(C)** Representative flow cytometry plots for intracellular cytokine staining of *in vitro*-differentiated Th1 and Th2 WT and IL31KO lymph node CD4 T cells. Histograms depict percent IL-31^+^ of live CD4+ cells. **(D)** Representative flow cytometry plots of IL-4 and IL-13 expression in *in vitro*-differentiated WT Th2 CD4 T cells (left panel all cells; right panel gated on IL-31^+^ cells as in C). Data shown are representative of 3 independent experiments (B and D). Datapoints reflect biological replicates, n=3 mice per genotype per condition. Error bars show mean +/-SD. Statistical significance was determined by unpaired 2-tailed student’s t-test. Only significant p-values are noted.

### *Il31* limits CD4 T cell cytokine production in response to topical HDM

To evaluate the contributions of endogenous *Il31 in vivo,* we turned to mouse models of dermatitis. We first examined a chronic house dust mite (HDM) model that involves twice-weekly dorsal neck skin barrier disruption followed by HDM ointment application (Fig. 2A)(*17, 18*). We assessed HDM-induced skin inflammation at week 5, a time point at which expression of *Il31* and its receptor subunits (*Il31ra* and *Osmr*) were detectable in treated skin (Fig. 2B and S2A) and scratching-induced skin injury was not yet appreciable. In both WT and IL31KO skin, histopathologic features of HDM-induced dermatitis included epidermal acanthosis, hypergranulosis, and mixed dermal infiltrates (Fig. 2C). HDM induced comparable expansion of cutaneous CD45^+^ hematopoietic cells including eosinophils, CD11b^+^Gr-1^+^ cells (neutrophils and monocytes), and CD4, CD8 and dermal *γδ* T cells in IL31KO and WT mice (Fig. 2D-E and S2B; see Fig. S3 for gating strategy). HDM treatment also triggered selective expansion and/or influx of cutaneous Foxp3^-^CD4^+^ T-effector cells as compared to Foxp3^+^CD4^+^ regulatory T cells (Tregs) (Fig. 2E).

**Fig. 2.**
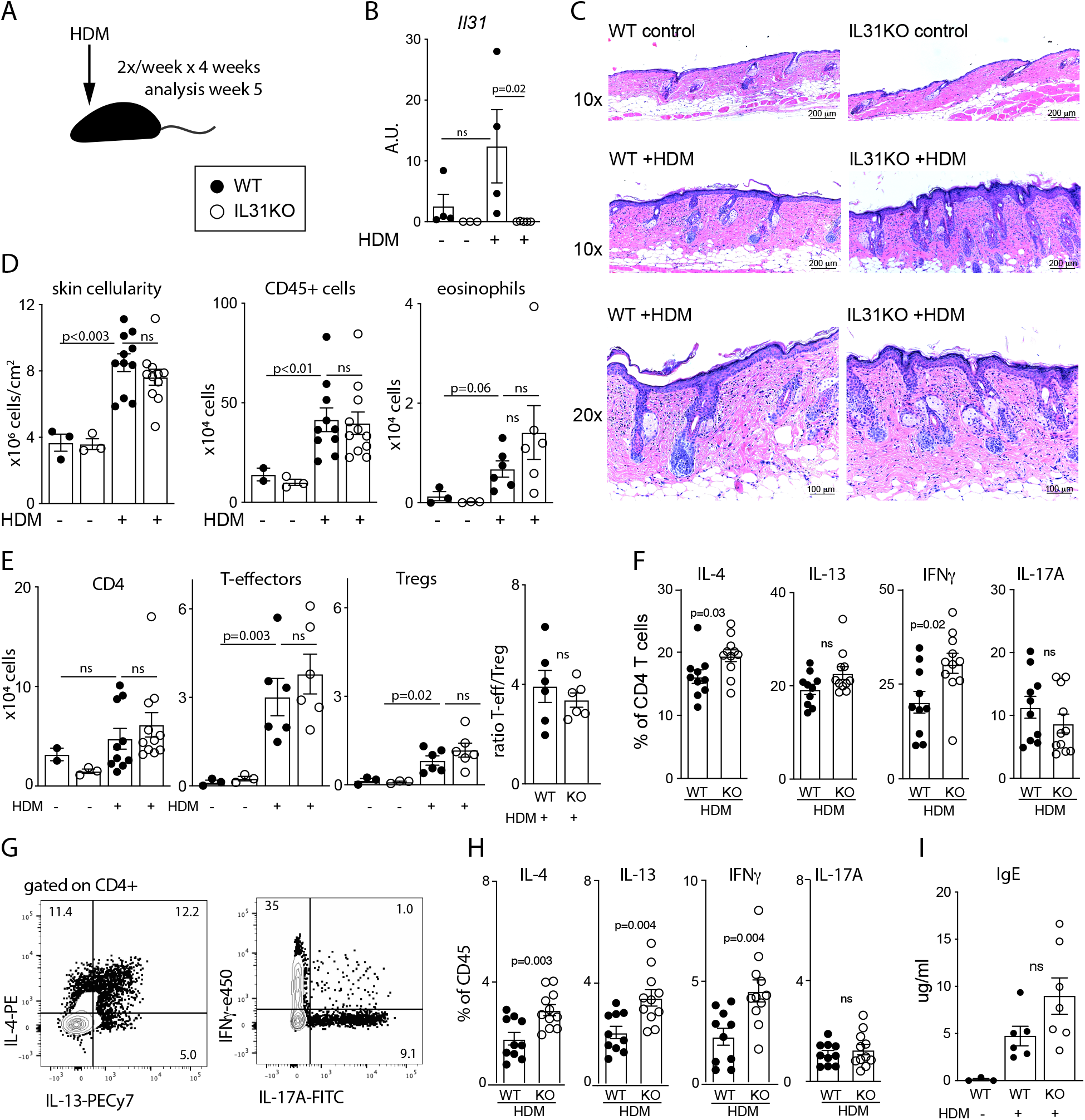
Increased cytokine-producing CD4 T cells distinguish dermal infiltrates in HDM-treated IL31KO animals. **(A)** Schematic of the 5-week HDM protocol used for skin histology and flow cytometry. **(B)** Relative abundance of *Il31* mRNA in total RNA extracted from dorsal neck skin of control and HDM-treated mice, measured by Taqman qPCR. A.U. arbitrary units. Representative of 2 experiments, n=3 mice per genotype and treatment condition. **(C)** Representative H&E-stained tissue sections from control or HDM-treated dorsal neck skin at 10x and 20x magnification. **(D)** Flow cytometry-based quantification of skin cellularity, CD45+ (hematopoietic) cells, and CD11b^+^SiglecF^+^ eosinophils. Cells shown per cm^2^ section of dorsal neck skin. Representative of 2 experiments, with n=3 controls and n≥6 for HDM-treated animals. See Fig. S3 for gating strategy. **(E)** Flow-cytometry-based quantification of CD4 T cell subsets as indicated. **(F)** Percent of HDM-treated skin CD4 T cells that express IL-4, IL-13, IFNγ, or IL-17A, as determined by flow cytometry after intracellular antibody staining. Pooled from 2 independent experiments including n=3 for controls and n≥6 mice for HDM. **(G)** Concatenated flow cytometry plots demonstrate gating for IL-4, IL-13 (left), IFNγ, IL-17A cytokine stains (right). **(H)** Cytokine expression data from **F** normalized by percent of CD45^+^ cells per sample. **(I)** ELISA-based quantification of serum IgE after 8 weeks of control-or HDM-treatment. All data points reflect unique biological replicates. Error bars displayed as mean +/-SD. Statistical significance was determined by unpaired 2-tailed student’s t-test. WT, black circles; IL31KO, open circles.

In analogous models of allergic airway inflammation induced by intranasal HDM administration, CD4 T cell-mediated inflammation is characterized by mixed Th1/Th2/Th17 infiltrates(*19*). Similarly, topical HDM-treatment of dorsal neck skin also resulted in expansion and/or recruitment of Th2, Th1 and Th17 cells in both WT and IL31KO mice (Fig. S2C). However, HDM-treated IL31KO mice generated significantly greater proportions of IFNγ and IL-4-producing CD4 T cells compared to WT controls (Fig. 2F; see Figs. 2G and S4 for gating strategy). When calculated as a percentage of CD45^+^ cells to correct for variation in skin CD4 T cell numbers in individual animals, IFNγ, IL-4, and IL-13-producing cells were all increased in HDM-treated IL31KO compared to WT (Fig. 2H). In control vehicle-treated animals of both genotypes, there were fewer CD4 T cells and therefore very few cytokine-producing T cells (Fig. S3C).

We also measured serum IL-4 and IgE in HDM-treated WT and IL31KO animals. Serum IL-4 was above the limit of detection in 5 of 12 treated IL31KO mice but in none of 13 treated WT mice (data not shown). This trend was reflected in the abundance of serum IgE, the production of which is dependent on IL-4. Serum IgE increased in HDM-treated WT animals compared to controls, and to an even greater degree in HDM-treated IL31KO animals (Fig. 2I). Overall, these results illustrate a role for IL-31 in restraining allergen-induced skin inflammation and IgE hyper-production.

### *Il31* also limits CD4 T cell IL-4 production in antigen-specific dermatitis

To determine whether the increased cytokine production we observed in HDM-treated IL31KO animals was reproducible in a T-cell antigen-dependent model of type 2 inflammation, we evaluated skin inflammation following sensitization and intradermal challenge with ovalbumin (OVA) (Fig. 3A).

**Fig. 3.**
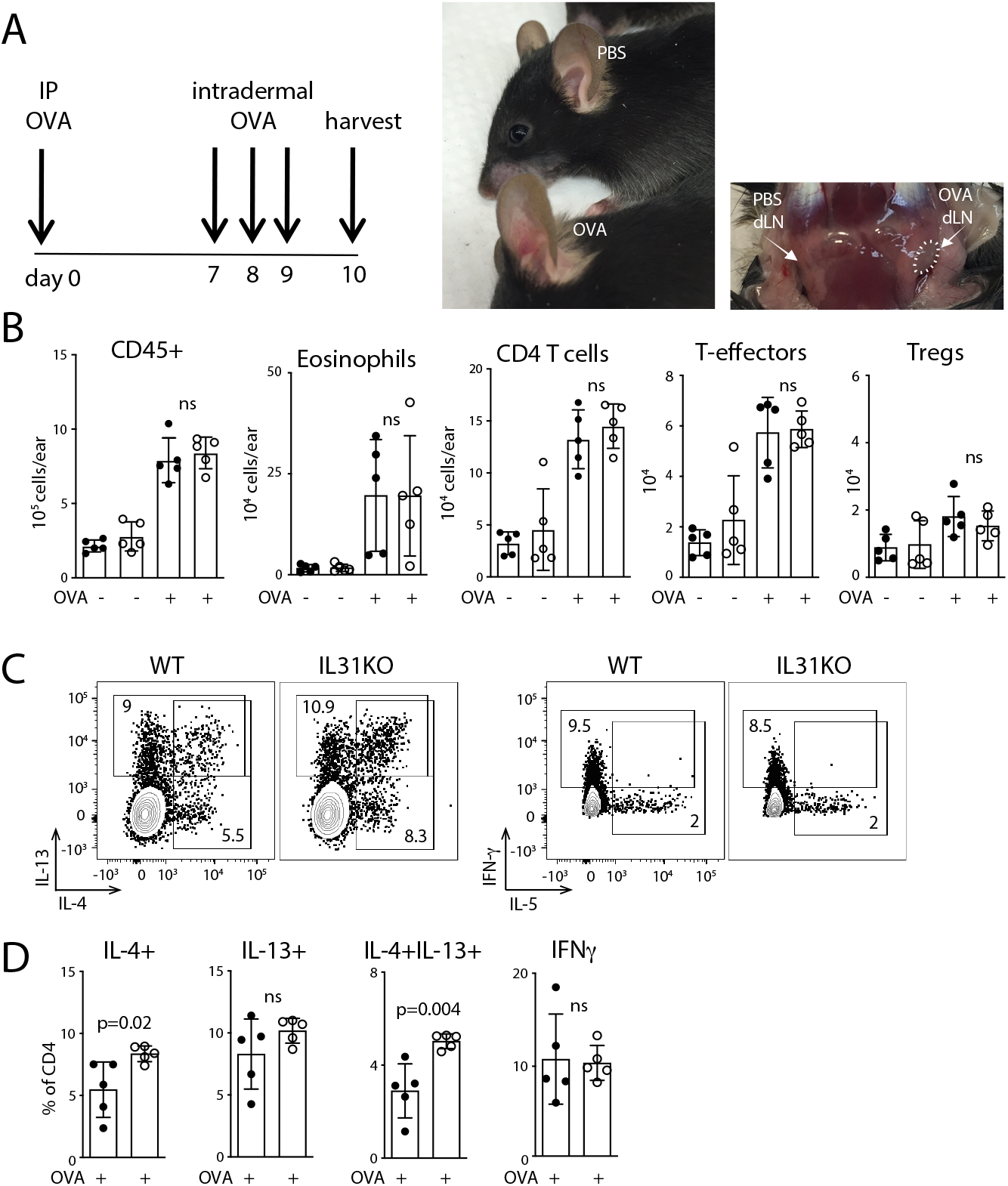
Increased Th2 cytokine production also observed in ovalbumin-challenged IL31KO animals. **(A)** Schematic of the 10-day ovalbumin intraperitoneal sensitization and ear skin challenge protocol. Photographs are representative of ovalbumin-treated and PBS control-treated ear skin and ear-draining lymph nodes (white arrows). **(B)** Flow cytometry-based enumeration of total skin CD45+ cells, eosinophils, and CD4 T cells (Foxp3-T effector and Foxp3+ T regulatory cells); data are displayed per ear. **(C)** Concatenated flow cytometry plots from ovalbumin-challenged ear skin, demonstrating gating after intracellular cytokine staining for IL-4, IL-13 (top row), IFNγ, IL-5 (bottom row). WT, left column; IL31KO, right column. **(D)** Percentage of OVA-challenged skin CD4+ T cells expressing cytokines as indicated. Data are representative of 3 independent experiments, including biological replicates (n=5 per condition), and displayed as mean+/-SD. Statistical significance was determined by unpaired 2-tailed student’s t-test. Gating strategy is analogous to the HDM model, as shown in Figs. S3 and S4. WT, black circles; IL31KO, open circles.

Intradermal OVA challenge resulted in expanded cutaneous populations of CD45^+^ hematopoietic cells, particularly eosinophils and both effector and regulatory CD4 T cells in both IL31KO and WT mice (Fig. 3B). However, as in HDM dermatitis, the proportion of IL-4-producing cutaneous CD4 T cells induced by OVA was increased in IL31KO compared to WT mice (Fig. 3C-D). The increase in IL-4 producing cells occurred within an IL-13 producing subset, so the overall effect of IL31KO was an increase in IL-4^+^IL-13^+^ double-producing CD4 T cells (Fig. 3D). From this result in a second model of dermatitis, we conclude that IL-31 participates in negative regulatory feedback of IL-4 production by cutaneous CD4^+^ effector T cells, supporting a general role for IL-31 in limiting allergen-induced adaptive type 2 responses in the skin.

Taken together, the IL31KO phenotype of increased IL-4, IL-13 and IFNγ cytokine production and increased serum IgE indicate that endogenous IL-31 attenuates rather than induces inflammation in skin. We considered the possibility that IL-31 acts in an autocrine fashion on CD4 T cells. However, we did not detect *Il31ra* expression in activated *in vitro*-differentiated Th2 cells (data not shown). These data suggest that IL-31 may be acting via cell populations that do express IL31RA, such as myeloid cells or neurons, to subsequently influence cytokine production in CD4 T cells.

### Disproportionate expansion of M2 macrophages in HDM-treated *Il31ra*-deficient skin

To characterize the effects of subtracting IL-31 signaling circuits from cutaneous myeloid populations, we performed single cell RNA-sequencing sequencing on sorted CD45^+^ cells from the skin of HDM-treated WT and *Il31ra*-deficient (IL31RAKO) mice. We resolved a combined total of 16284 cells, 47% lymphoid (7509) and 53% (8775) myeloid (Fig. 4A). Within the myeloid compartment, we observed a 1.72-fold enrichment of IL31RAKO cells (5549 KO, 3226 WT), consistent with the exacerbated cutaneous inflammatory response to topical HDM observed in IL31KO animals by flow cytometry.

**Fig. 4.**
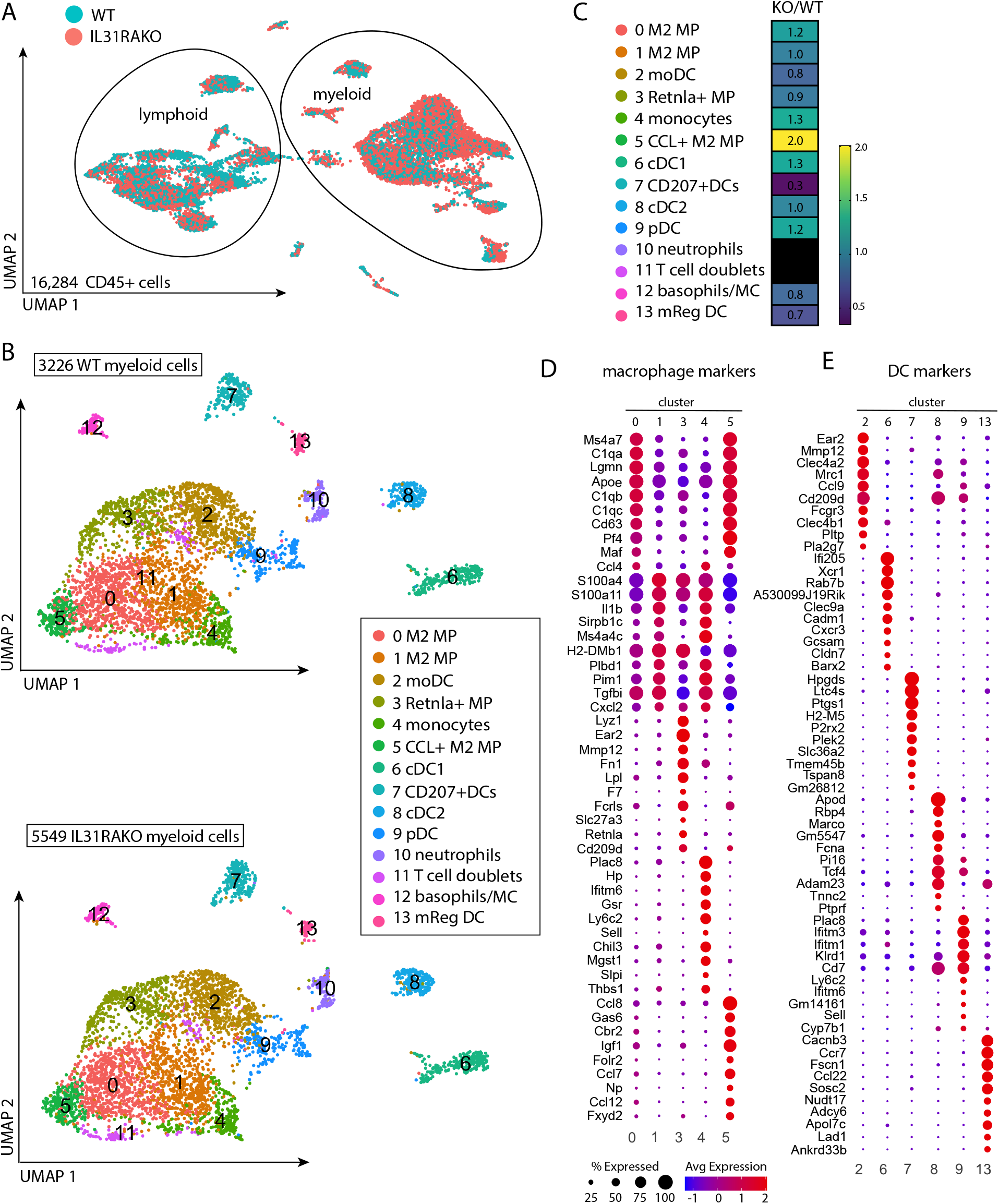
Selective expansion of cutaneous M2 macrophage subsets in HDM-treated IL31RAKO mice. **(A)** UMAP plot of merged WT and IL31RAKO scRNA-seq files (combined total 16,284 cells) demonstrates overall clustering by lineage (lymphoid vs myeloid). Individual cells are displayed by genotype using the Seurat shuffle command. Turquoise = WT cells; salmon = IL31RAKO cells). IL31RAKO and WT scRNA-seq files were each generated from sorted CD45+ cells pooled from n=4 animals. **(B)** UMAP plot after subclustering the myeloid cells indicated in A. Top: WT cells; Bottom: IL31RAKO cells. Cell identities of the 13 myeloid cell clusters (inset box) were determined by inspection of cell marker genes identified by differential gene expression (DEG) analysis (see Fig. S5). **(C)** Heat map depicting proportional representation of IL31KO and WT cells in each myeloid cluster, where numbers reflect the ratio of Il31RAKO to WT cells per cluster after normalization to correct for the 1.72-fold greater number of total cells resolved in the IL31RAKO sequencing file. **(D)** Dot plots depict expression of the top 10 DEGs for each macrophage cluster mapped across all macrophage clusters (clusters 0,1,3,4,5). **(E)** Dot plots depict expression of the top 10 DEGs for each dendritic cell cluster (clusters 2,6,7,8,9,13) mapped across all DC clusters. As indicated by the legend, the diameter of each circle indicates the proportion of cells in that cluster that express the gene, and the heat map indicates average expression per cell in that cluster by read count.

Unsupervised subclustering analysis of the 8775 myeloid cells identified 13 distinct populations, including 5 monocyte-macrophage clusters, 6 dendritic cell subsets, and a co-cluster of mast cells and basophils (Fig. 4B; see Fig. S5 for cluster marker genes). IL31RAKO and WT cells were represented in all 13 clusters, but IL31RAKO cells were overrepresented in all but one: cluster 7 (*Cd207*^+^ DCs). To compare proportional representation of IL31RAKO and WT cells per cluster, we normalized to the overall ratio of IL31RAKO to WT myeloid cells (a factor of 1.72) (Fig. 4C). After normalization, IL31RAKO cells were still overrepresented in multiple myeloid clusters, including two large macrophage populations (cluster 5, 2.0-fold KO>WT and cluster 0, 1.2-fold KO>WT) that together account for 2,101 cells, or 23.9% of total myeloid cells sequenced. We only resolved *Il31ra* and *Osmr* transcripts on rare cells (Fig. S6A). Oncostatin M (*Osm)* was expressed in cells of multiple clusters, with no appreciable differences between IL31RAKO and WT (Fig. S6B).

We used differentially expressed gene (DEG) analysis to define cluster markers for monocyte-macrophage and dendritic cell subsets (Fig. 4D-E). The macrophage subsets specifically enriched in KO skin (clusters 0 and 5) were highly similar to one another, and distinct from other macrophage clusters (Fig. 4D). Genes whose selective expression drove this clustering included *Ms4a7*, *Apoe*, *Pf4* (Cxcl4), and *Maf* (Figs. 4D and 5A). Cluster 0 and 5 cells also expressed high levels of complement receptors *C1qa*, *C1qb,* and *C1qc*, reflective of IL-4-driven type 2 macrophage (M2) differentiation and increased phagocytic capacity(*20*). Additional cluster 0 and 5 markers were enriched in other macrophages (clusters 1 and 3) and monocytes (cluster 4), but at lower read counts and in smaller percentages of cells (Fig 4D and 5A).

**Fig. 5.**
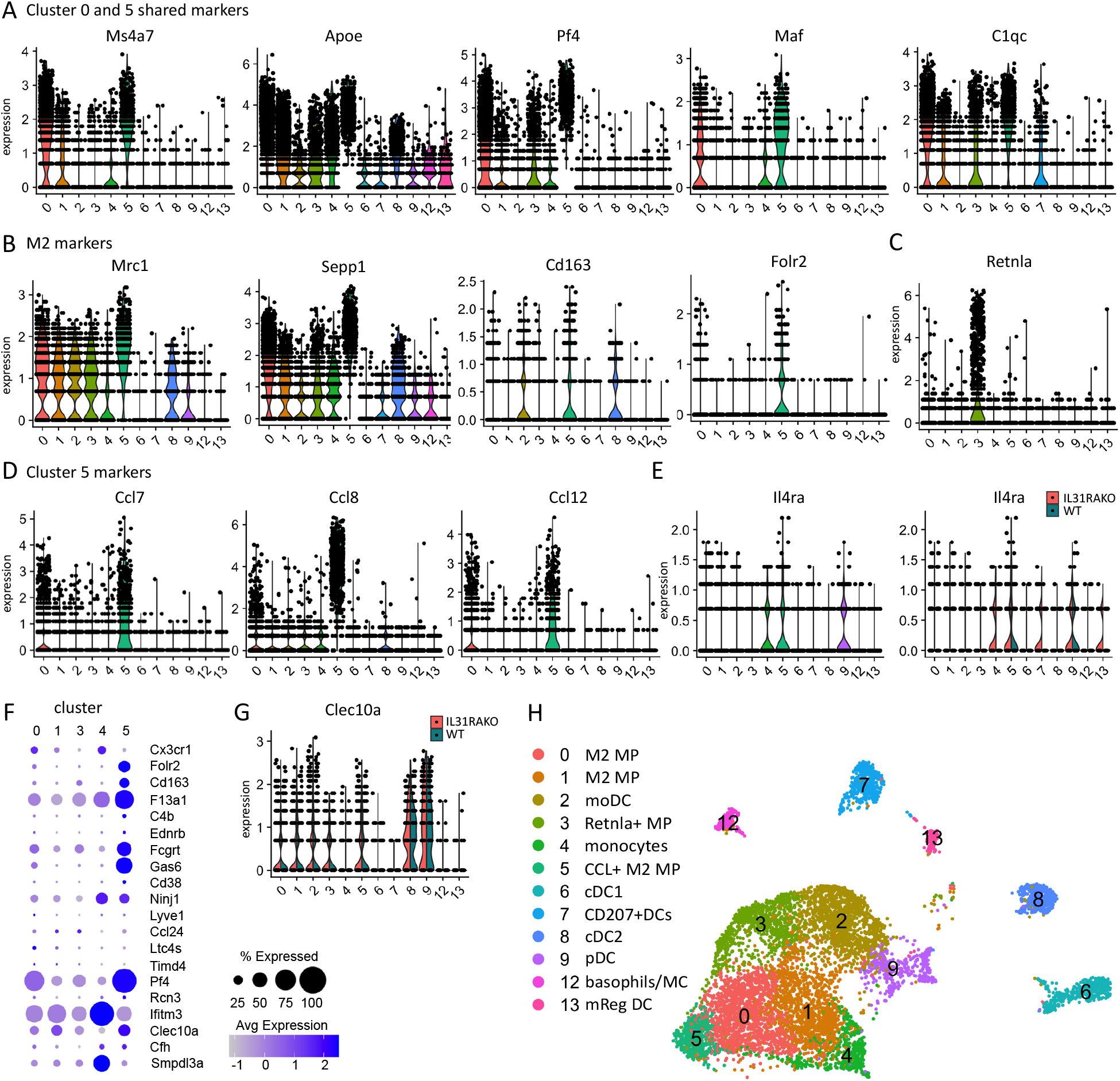
Macrophage subsets enriched in HDM-treated IL31RAKO skin are distinguished by transcripts encoding M2 markers, CCR2 ligands and IL4RA. **(A)** Expression of shared DEG markers for cluster 0 and cluster 5, displayed as violin plots. Y-axis indicates normalized gene expression by read count. **(B)** Violin plots depicting expression of M2 markers *Mrc1, Sepp1, Cd163,* and *Folr2* mapped across myeloid clusters demonstrate enriched expression in multiple macrophage clusters, including 0 and 5. **(C)** Violin plot depicting expression of M2 marker gene *Rentla* (Fizz1). **(D)** Additional cluster 5 markers include *Ccl7*, *Ccl8*, and *Ccl12*, CCR2 ligands encoded in a chemokine cluster on mouse chromosome 11. **(E)** Violin plot (left) highlights expression of *Il4ra* in cluster 5 (CCL+ macrophages) and cluster 9 (pDCs). Split violin plots (right) reveal additional *Il4ra* expression in IL31RAKO cluster 7 (Cd207+DCs) and cluster 13 (mReg DCs) cells. **(F)** Dot plot depicting expression of markers for Cx3cr1^low^ tissue macrophages* in our scRNAseq macrophage clusters. *Genelist extracted from Chakarov et al, Table S3(*24*)). **(G)** Violin plot depicting expression of HDM-response factor *Clec10a* in our dataset, split by sample (Turquoise = WT cells; salmon = IL31RAKO cells **(H)** Cell identities by cluster, with color scheme. [Note: Cluster 10 (neutrophils) and cluster 11 (T cell-myeloid doublets) were excluded from this analysis, so the color scheme differs from Figure 4 UMAP plots. Cluster numbers are consistent.]

We next examined macrophage subsets for expression of M2 differentiation markers. Transcripts encoding the canonical M2 marker *Arg1* were rare (Fig. S6C), and the M1 marker *Nos2* was undetectable. However, multiple macrophage clusters contained a large proportion of cells that expressed *Sepp1* and *Mrc1* (mannose receptor), genes also associated with M2 macrophage differentiation(*21*) (Fig. 5B), suggesting that most macrophages in HDM-treated skin adopt an M2 fate. Nearly all cluster 5 macrophages expressed abundant *Mrc1* and *Sepp1* and some were enriched for additional M2 markers *Folr2* and *Cd163* (hemoglobin scavenger receptor)(*21*) (Fig. 5B). *Retnla* (Fizz1) was enriched in cluster 3 macrophages (Fig. 5C), which were equally represented in IL31RAKO and WT samples (Fig. 4C). Altogether, these scRNA-seq results highlight heterogeneous subsets of M2 macrophages in HDM-inflamed skin, some limited by IL-31.

Cluster 5-specific DEG markers (Fig. 5D; see also 4C and S6D) included Mertk ligand *Gas6,* and *Ccl7*, *Ccl8*, and *Ccl12*. The latter encode monocyte chemoattractant ligands for CCR2, a chemokine receptor broadly expressed by monocytes, macrophages and monocyte-derived dendritic cells (Fig. S6E). Based upon the two-fold enrichment of cluster 5 *Ccl7*^+^*Ccl8*^+^*Ccl12*^+^ macrophages in IL31RAKO skin, we conclude that the lack of IL-31 signaling in IL31RAKO mice enables a feedforward loop whereby M2-polarized cutaneous macrophage populations expand in response to HDM by recruiting CCR2^+^ monocyte or macrophage precursors (cluster 4, 1.3-fold) from peripheral blood.

Although *Il4ra* was not defined as a DEG for any cluster, cluster 5 cells did exhibit the highest expression of *Il4ra* (Fig. 5E). *Il4ra* expression was also detected in plasmacytoid DCs (cluster 9). Notably, in IL31RAKO but not WT cells we detected additional *Il4ra* expression in cluster 4 monocytes, cluster 7 Cd207+DCs, and cluster 13 mReg DCs, suggestive of increased IL-4 responsiveness in the myeloid compartment in the absence of IL-31/IL31RA signaling.

Cx3cr1 abundance distinguishes dermal-resident macrophages (Cx3cr1^low^) from Cx3cr1^high^ macrophages that interact with sensory neurons and continuously repopulate from bone marrow-derived peripheral blood monocyte precursors(*22–24*). In HDM-treated skin, expression of *Cx3cr1* mRNA was too similar across macrophage subsets to allow us to define clusters using this marker alone (Fig. S6F). However, based upon common expression of DEGs *Maf*, *Mrc1*, *Sepp1*, and *Cd163*, we hypothesize that clusters 0 and/or 5 most closely approximate tissue-resident Cx3cr1^low^ macrophages. Mapping Cx3Cr1^low^ macrophage signature genes(*24*) onto our dataset (Fig. 5F) supports the conclusion that cluster 5 macrophages, which are overrepresented by 2-fold in IL31RAKO compared with WT skin, are the tissue-resident subset.

The effects of IL31RA deficiency on dendritic cell (DC) subsets in skin included modest enrichment of IL-4-responsive plasmacytoid DCs (cluster 9, 1.2-fold) and *Irf8*^+^ cDC1s (cluster 6, 1.3-fold) (Fig. 4C). In addition, fewer mRegDCs(*25*) (cluster 13, 0.7-fold) were detected in IL31RAKO skin. The proportion of *Cd301b*^+^ *Irf4*^+^ cDC2s (cluster 8) was unaffected by IL31RA deficiency. Cluster 8 cDC2s and cluster 9 pDCs were the only DCs to express HDM response factor *Clec10a* (Fig. 5G)(*26*).

Overall, this comparative scRNA-seq analysis of IL31RAKO and WT HDM-treated skin infiltrates revealed a role for IL31RA signaling in controlling myeloid inflammation in allergen-exposed skin, in particular by limiting IL-4-responsive M2 macrophage subsets. We suggest that one of these control mechanisms may be through IL-31-dependent signals limiting a chemokine-dependent feedforward loop that otherwise perpetuates expansion and differentiation of cutaneous allergen-induced IL-4-responsive myeloid cells.

### Decreased spontaneous scratching in HDM-treated IL31KO reveals a selective sensory deficit

Skin-innervating IL31RA^+^ pruritoceptors constitute the major non-hematopoietic population of IL-31 responsive cells. To determine whether IL31RA^+^ neurons are necessary contributors to allergen-induced pruritoception, we used the Lamotte method(*27*) to evaluate behaviors attributable to grooming (frontpaw wiping) versus itching (hindpaw scratching) in the chronic HDM allergen model (Fig. 6A).

**Fig. 6.**
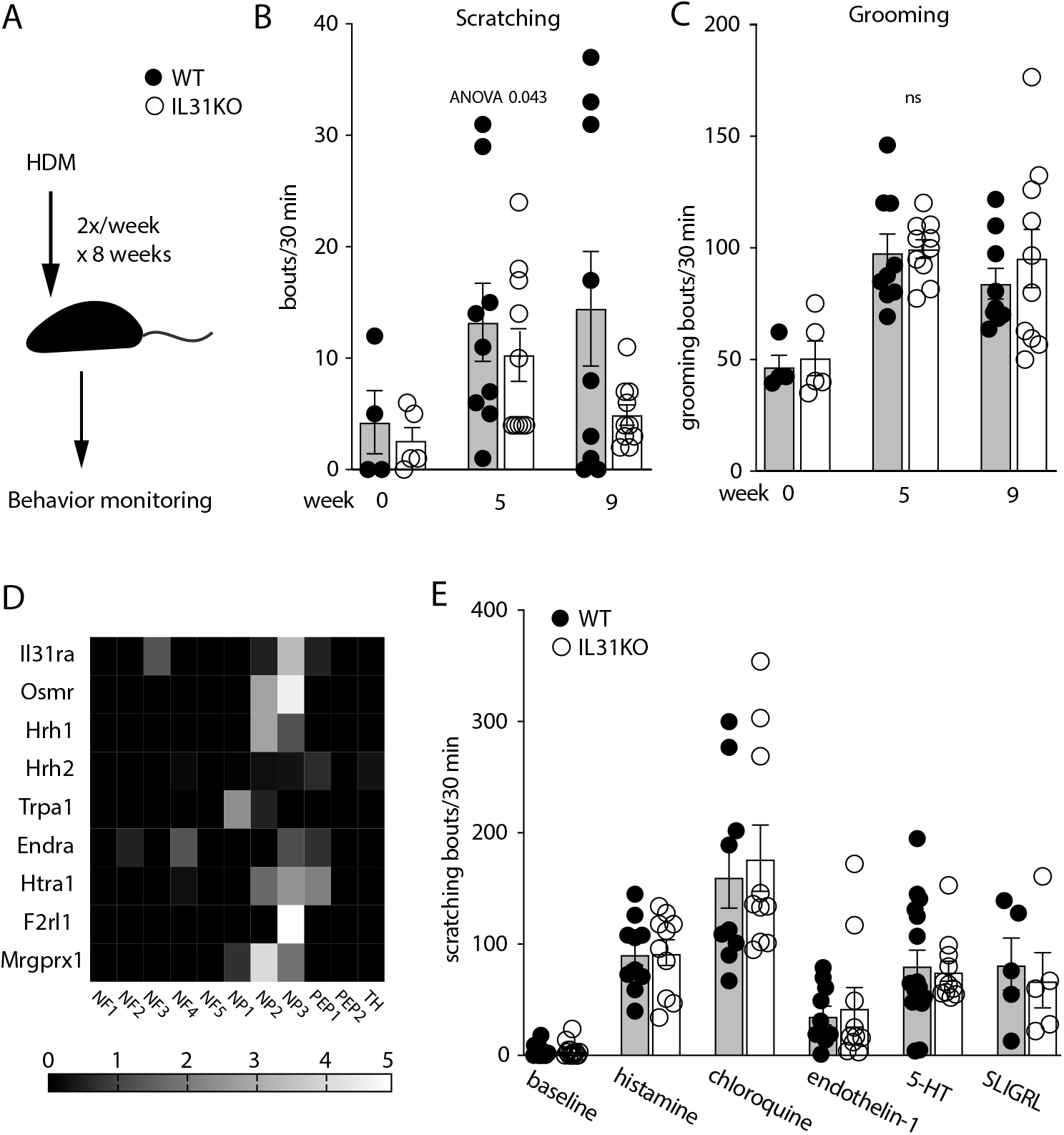
HDM-sensitized IL31KO mice display a deficit in spontaneous scratching. **(A)** Schematic of the 8-week chronic HDM epicutaneous sensitization strategy; behavior monitoring was performed in week 9. **(B)** Spontaneous scratching bouts (hindpaw) per 30 minute window, measured serially in a cohort of WT and IL31KO animals at week 0 (before treatment), week 5, and week 9 (after completion of treatment). **(C)** Spontaneous grooming bouts (forepaw) per 30 minutes, measured in the same videos as B. **(D)** Heat map depicts gene expression of select pruritogen receptors in dorsal root ganglia afferent neuron subsets, extracted from Usoskin et al(*29*). **(E)** Scratching bouts per 30 minute window, assessed immediately following injection of synthetic pruritogens as indicated. Black circles, WT; open circles, IL31KO. Data are pooled from 2 independent experiments, with n ≥4 mice per genotype (B and C) or n≥5 animals per genotype (E). Each dot represents an individual mouse. Statistical significance of the difference between WT and IL31KO was determined by Welch’s one-way ANOVA across all timepoints (B and C) or by unpaired 2-tailed student’s t-test (E). Error bars indicate mean +/-SD.

Baseline grooming and scratching did not differ between IL31KO and WT mice (Fig. 6B-C). In WT animals, HDM treatment induced a modest increase in spontaneous scratching frequency at week 5 and the majority of mice continued to scratch through week 9, consistent with previous reports(*28*). IL31KO animals also displayed increased spontaneous scratching at week 5, but by week 9 the frequency of scratching in these mice fell to near-baseline (Fig. 6B). In contrast, grooming frequency of IL31KO and WT mice tracked together throughout the HDM-treatment period (Fig. 6C), increasing similarly in HDM-treated animals of both genotypes.

We examined published DRG neuron scRNA-seq data for patterns of pruritogen receptor expression that could impact HDM-induced scratching in IL31KO mice. The NP3 subset of DRG sensory afferents are notable for expression of *Il31ra and Osmr* and additional pruritogen receptors PAR2 (encoded by *F2rl1*) and Mrgprc11 (encoded by *Mrgprx1*) (Fig. 6D, excerpted from Usoskin et al(*29*)). As transgenic overexpression of *F2rl1* exacerbates HDM-induced scratching and leads to upregulation of multiple pruritogen receptors in trigeminal ganglia (TG) in the setting of HDM-induced dermatitis, including *Il31ra* and *Osmr(30, 31),(28)*, co-expression of *Il31ra*, *Osmr* and *F2rl1* in NP3 neurons may have functional consequences for allergic inflammation-associated pruritus. Additional allergy-associated pruritogen receptors (*Il4ra, Hrh1* and *Hrh2, Endra*, and *Htra1*) are more broadly expressed on DRG sensory neuron subsets, and no particular relationship to the IL-31 receptor subunits was apparent (Fig. 6D).

To evaluate potential *Il31*-dependent perturbation of itch pathways mediated by co-expressed pruritogen receptors, we also monitored scratching frequency in response to acute subcutaneous injection of a panel of pruritogens, including histamine, endothelin-1, serotonin (5-HT), and SLIGRL, a PAR2 ligand. IL31KO mice scratched at frequencies indistinguishable from WT in response to these pruritogens (Fig. 6E), indicating that these ligands function independently of IL-31. In addition, these data suggest that attenuated scratching in IL31KO mice in the chronic HDM allergen model does not depend on these other allergy-associated pruritogen receptors, and may be attributed to altered IL31RA/OSMRβ signaling.

### IL-31 restrains expression and activation of PAR2 in sensory neurons

Given the decreased scratching observed in HDM-treated IL31KO mice, we hypothesized that *Il31-*deficiency may alter the proportion of sensory neurons that express *Il31ra* or *F2rl1*(PAR2). To test this hypothesis, we used *in situ* hybridization to monitor expression of *Il31ra* and *F2rl1* in TG neurons from mice exposed to HDM on cheek skin. *Il31ra* and *F2rl1* transcripts were expressed in overlapping subsets of *Trpv1*^+^ small-diameter neurons (Fig. 7A), consistent with published DRG scRNA-seq data (Fig. 6D) (*29*)). These were largely distinct from *Il4ra^+^* TG neurons (Fig. 7B), as previously described(*28*). *Il31ra* and *F2rl1* transcription also overlapped in *Trpv1*^+^ neurons in IL31KO; in the IL31KO there were significantly more *Il31ra*^+^, *F2rl1*^+^, and *Il31ra*^+^*F2rl1*^+^ TG neurons than in WT (Fig. 7C). HDM treatment also increased the number of neurons expressing these pruritoceptors in both WT and IL31KO TG, but only the increase in *F2rl1*^+^ neurons in IL31KO TG was significant post-HDM treatment (Fig. 7C). This unexpected result indicates that intact IL-31 signaling limits neuronal expression of PAR2 and therefore may limit pruritoceptor activation by PAR2 ligands.

**Fig. 7.**
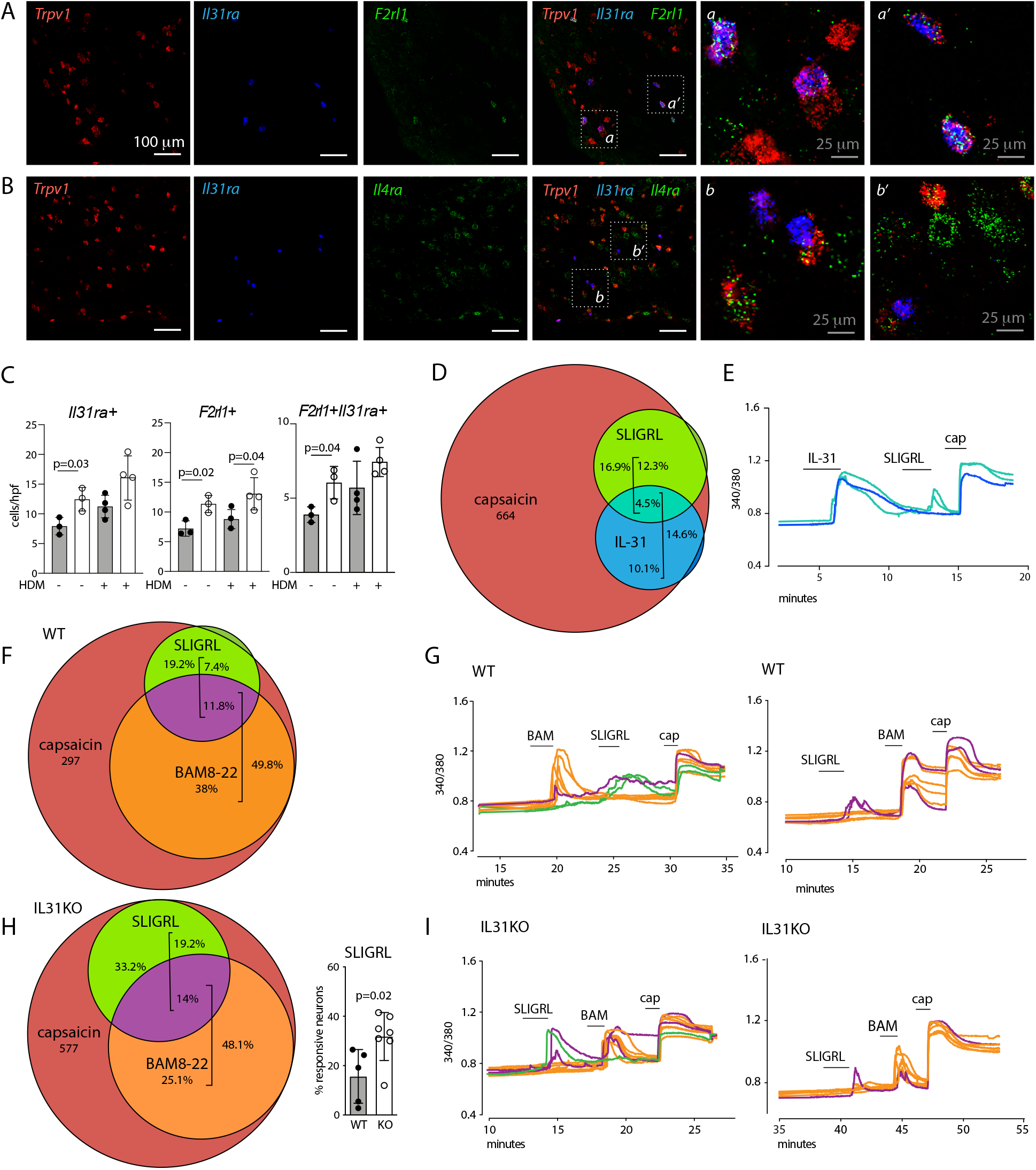
IL31KO mice have an increased proportion of sensory neurons responsive to PAR2 ligand SLIGRL. **(A)** Representative images of *Trpv1*, *Il31ra*, and *F2rl1 in situ* hybridization in trigeminal ganglia (TG) tissue sections from untreated animals. **(B)** Representative images of *Trpv1*, *Il31ra*, and *Il4ra* by *in situ* hybridization in TG tissue sections from untreated animals. **(C)** Number of TG cells per high power field (hpf) stained with probes for *Il31ra*, *F2rl1* or both. **(D)** Venn diagram indicating the proportion of WT dorsal root ganglion (DRG) neurons responding to each of the indicated ligands (capsaicin, SLIGRL, and recombinant IL-31). Capsaicin response marks TRPV1^+^ neurons. **(E)** Representative 340/380 traces from calcium imaging of Fura-2 loaded lumbar DRG neurons that were monoresponsive to IL-31 (blue) or double-responsive to IL-31 and SLIGRL (turquoise). **(F)** Venn diagram indicating the proportion of WT DRG neurons responding to each of the indicated ligands (capsaicin, SLIGR, and BAM8-22). **(G)** Representative 340/380 traces from Fura-2-labeled WT lumbar DRG neurons monoresponsive to BAM8-22 (orange), monoresponsive to SLIGRL (green), or double-responsive to BAM8-22 and SLIGRL (purple). **(H)** Venn diagram indicating the proportion of IL31KO DRG neurons responding to the indicated ligands (capsaicin, SLIGRL, and BAM8-22). Histogram depicts SLIRGL-responsive WT and IL31KO neurons as a percent of capsaicin-responsive (TRPV1^+^) neurons, with each dot representing an individual mouse. **(I)** Representative 340/380 traces from calcium imaging of Fura-2-labeled IL31KO lumbar DRG neurons that were monoresponsive to BAM8-22 (orange), monoresponsive to SLIGRL (green), or double-responsive to BAM8-22 and SLIGRL (purple). Black circles, WT; open circles, IL31KO. Numbers indicate total responding neurons recorded from 10 WT mice (D); 5 WT mice (F); or 7 IL31KO mice (H). Significance was determined by unpaired 2-tailed student’s t-test (C and H).

Finally, we performed calcium imaging of *ex vivo* lumbar DRG neurons, and confirmed the pattern of IL31RA/PAR2 co-expression at a functional level. We recorded a total of 1625 dissociated DRG neurons from WT C57Bl/6 animals (Fig. 7D-E). Of these, 664 (40.9%) responded to the TRPV1 agonist capsaicin, 102 (6.3%) responded to recombinant IL-31, and 112 (10.5%) responded to PAR2 ligand SLIGRL. The vast majority of IL-31-responsive (95%) and SLIGRL-responsive (94.6%) DRG neurons were also capsaicin-responsive, consistent with prior reports and with our *in situ* hybridization analyses of the distribution of their receptors, IL31RA and PAR2, in TRPV1^+^ primary afferent DRG neurons (*29, 32*). Within the capsaicin-responsive subset, 16.9% were SLIGRL-responsive, 14.6% were IL-31-responsive, and 4.5% responded to both SLIGRL and IL-31, consistent with prior reports(*32*).

To further test whether *Il31*-deficiency alters the frequency of PAR2^+^ neurons, we compared WT and IL31KO lumbar DRG neuron responses to SLIGRL. Among capsaicin-responsive DRG neurons, 19.2% were SLIGRL-responsive in WT DRG (Fig. 7F-G) compared to 33.2% in IL31KO DRG (Fig. 7H-I). This difference was significant when normalized to the number of capsaicin-responsive DRG neurons per mouse (Fig. 7H histogram). However, an increased percentage of SLIGRL-responsive cells is insufficient to affirm increased PAR2 activation in IL31KO because SLIGRL can also activate a second pruritogen receptor, Mrgprc11(*33*). Calcium flux in response to the selective Mrgprc11 ligand Bam8-22(*34*) was comparable within the set of capsaicin-responsive DRG neurons in WT (49.8%; Fig. 7F-G) and IL31KO (48.1%; Fig. 7H-I). As only the SLIGRL-responsive Bam8-22-nonresponsive fraction differed between genotypes (7.4% in WT vs. 19.2% in KO), we conclude that the observed increase in SLIGRL-responsive neurons is secondary to an increase in PAR2 expressing-neurons, and unrelated to Mrgprc11. Furthermore, the increased proportion of PAR2^+^ SLIGRL-responsive DRG neurons in IL31KO mice suggests that a net decrease in HDM-induced scratching in IL31KO occurs independently of PAR2, and underscores the necessary contribution of IL-31:IL31RA-mediated pruritoceptor activation to HDM-induced itch.

## DISCUSSION

We demonstrated significant effects of *Il31*-deficiency on adaptive type 2 cytokine production in two discrete models of allergic skin inflammation, and uncovered a role for IL-31 in regulating the composition of allergen-induced IL-4-responsive type 2 myeloid cells in skin. These findings indicate that, when present, IL-31 limits cutaneous and systemic type 2 inflammatory responses. We also demonstrated that IL-31 influences itch-provoking sensory neurons during HDM-induced dermatitis. The consequence of *Il31*-deficiency is a net deficit in environmental allergen-induced itch despite intact behavioral indices of itch in response to other common allergy-associated pruritogens. Taken together, we conclude that IL-31 modulates non-redundant pathways that contribute to the development of both cutaneous inflammation and environmental allergen-induced pruritus.

Other cytokines have the capacity to communicate directly with both cutaneous inflammatory cells and sensory neurons(*35*), (*36*). However, these cytokines (epithelial alarmin TSLP and canonical type 2 cytokine IL-4) enhance both skin inflammation and sensory responses. IL-31, on the other hand, serves opposing roles. While it limits allergic inflammation, it enhances pruritoceptive responses. In sum, IL-31 functionally uncouples itch and allergic inflammation, the first cytokine known to do so.

### IL-31 and sensory neurons

Both gain-and loss-of-function experiments have demonstrated the pruritogenic properties of IL-31(*5, 6, 32*). Here, we tested the contribution of IL-31 in a complex, clinically-relevant tissue microenvironment: cutaneous inflammation stimulated by chronic HDM exposure. Numerous exogenous and endogenous pruritogens characterize HDM-treated skin. Therefore, we hypothesized that the reduced HDM-induced scratching observed in *Il31*-deficient animals could reflect secondary effects of altered (reduced) expression or activation of co-expressed pruritogen receptors. To address this possibility we focused on PAR2 because HDM-encoded proteases are capable of activating protease-activated receptors(*37*),(*38*) and PAR2 transgenic animals are hypersensitive to HDM-induced pruritus(*28, 30*). Unexpectedly, in IL31KO mice we observed a net increase in the number of PAR2^+^ (SLIGRL monoresponsive) neurons and IL31RA^+^PAR2^+^ (IL-31 and SLIGRL double-responsive) neurons. This could have conferred increased neural responsiveness to HDM, but it did not. That an *Il31*-dependent decrease in HDM-associated spontaneous scratching occurred despite increased numbers of PAR2^+^ pruritoceptors indicates that these co-expressed pruritogen receptors function independently within neurons, and underscores the non-redundant contribution of IL-31 to allergen-associated itch pathways.

### IL-31 and cutaneous myeloid cells

HDM treatment also uncovered *Il31-*dependent regulation of type 2 cytokine production by skin T cells, as well as monocyte-macrophage chemoattraction and M2 macrophage polarization. Release of mediators from the peripheral terminals of sensory neurons can influence cutaneous myeloid populations(*39*). However, as IL31RA is expressed at several stages of myeloid cell differentiation (*1, 40*), *Il31*-dependent effects are also likely to be mediated by hematopoietic cell-intrinsic circuits.

ScRNA-seq revealed an *Il31ra*-dependent expansion of skin-infiltrating myeloid cells, with disproportionate increases in highly-phagocytic and chemokine-producing macrophage subsets. Although these macrophage subsets (clusters 0, 1 and 5) did not express transcripts for the canonical markers of either M1 or M2 fates, DEGs enriched in these cells suggest that they are indeed M2 macrophages. Analogous M2 macrophages that express *Sepp1, Mrc1*, *Cd163*, and *Cbr2* but not *Arg1* have also been described in the setting of atherosclerosis(*21*), and *Folr2^+^* M2 macrophages were described in the oxazolone model of dermatitis(*23, 41, 42*).

Two myeloid cell populations have been associated with the cutaneous response to HDM: CX3CR1^high^ macrophages and cDC2s(*22, 24, 43*). However, both populations were equally represented in IL31RAKO and WT HDM-treated skin, suggesting that IL-31/IL31RA-dependent skewing of the cutaneous type 2 inflammatory response does not track with these cells.

### IL-31-dependent regulation of type 2 inflammation

Taken together, our studies advance the current understanding of IL-31 signaling in cutaneous type 2 inflammation by providing support for IL-31 as a negative immunoregulator. This possibility was previously suggested by studies demonstrating that *Il31ra*-deficient animals develop exacerbated inflammatory responses in colitis and allergic airway models. Interpretation of those studies is complicated by the possibility that phenotypes were driven by increased oncostatin M signaling secondary to altered OSMRβ availability in the absence of IL31RA(*10*). By genetically deleting the *Il31* cytokine rather than the *Il31ra* receptor subunit, our *Il31*-deficient animal experiments bypass that caveat.

Independently-derived *Il31*-deficient mice also exhibited decreased scratching in contact dermatitis models(*5*). Although no significant alterations in serum cytokine production were reported, IFNγ and TNF*α* production trended toward increases in those *Il31*-deficient mice, consistent with results presented here. Differences in the magnitude of our findings likely reflect the acute versus chronic nature of cutaneous allergen exposure and sensitivity differences between cytokine detection techniques. Here, cytokine detection by intracellular antibody staining for flow cytometry allowed us to evaluate CD4 T cell cytokine production at both single cell and population perspectives.

### Limitations

Unfortunately, *Il31ra* and *Osmr* transcripts were too rare in our scRNA-seq dataset to allow us to identify direct IL-31-responsive myeloid population(s). Further studies in lineage-specific *Il31ra* conditional mutant mice are needed to determine whether the IL-31-responsive IL31RA^+^ cells responsible for limiting type 2 inflammatory responses are myeloid or neural. From there, it will be exciting to evaluate the extent to which the *Il31*-dependent regulatory circuits we identified here, in mouse models, also participate in IL-31-mediated regulation of type 2 tissue inflammatory responses in atopy-prone patients as well.

Taken together, our data provide the strongest evidence to-date that IL-31 has dual functions as a pruritogen and immunoregulatory cytokine. Given the significant translational relevance of these conclusions to clinical dermatology, we note similar results from anti-IL31RA (Nemolizumab) clinical trials for AD(*44–46*). Nemolizumab-treated AD patients experienced rapid improvement in pruritus but delayed improvements in dermatitis metrics, and paradoxical dermatitis flares (19-21% vs 13% placebo in phase IIB(*47*); 23-24% vs 21% placebo in phase III(*44*)). The phase III study also uncovered “abnormal cytokine” levels (7% treatment vs 0% placebo), and elevated serum levels of the IL-4-responsive epithelial cytokine TARC, with an ∼3-fold mean increase from baseline in all Nemolizumab-treated AD patients (*44, 46*). Overall impressive clinical improvement in Nemolizumab-treated AD patients despite these increased inflammatory markers in a sizable subgroup suggests functional dominance of IL31RA^+^ pruritoceptive pathways in this disease. We interpret these clinical data as consistent with the complex phenotypic profile of our *Il31*-deficient animals, in whom pruritus and dermatitis pathways are similarly decoupled.

## MATERIALS AND METHODS

### Study Design

The objective of this study was to elucidate the effects of IL-31 signaling on the intersecting processes of cutaneous inflammation and itch, which are both active during allergic and atopic skin inflammation in genetically-predisposed individuals. To do so, we employed an established mouse model for allergic dermatitis induced by serial exposure to house dust mite, a common environmental allergen. As readouts for *Il31*-and *Il31ra*-dependent differences in cutaneous inflammation, we used a combination of flow cytometry and scRNA-seq of skin infiltrates. To detect alterations in sensory neuron responses, we performed spontaneous and provoked behavior assays, dorsal root ganglion *in situ* hybridization, and calcium imaging. Details of these experimental methods are delineated in this section.

### 31KO transgene design and transgenic animal screening

The *IL31^tm1e(EUCOMM)Wtsi^* transgene allele interrupts the endogenous *Il31* locus (mm10 ch5:123480419-123482049, minus strand) at position 123481232; it was generated by the European Mutant Mouse Consortium (EUCOMM) according to the insertional mutagenesis scheme described by Skarnes et al(*15*). The *IL31^tm1e(EUCOMM)Wtsi^* transgene contains a neomycin-selection cassette flanked by FRT sites, and paired loxp sites flanking *Il31* exon 2. When excised by serial genetic crosses to Flp-recombinase-and *β*-actin-Cre recombinase-expressing mouse strains, the resulting *IL31^null^* allele causes a frameshift mutation and generation of a nonsense transcript.

C57Bl/6 frozen embryos containing the *IL31^tm1a(EUCOMM)Wtsi^* transgene were obtained from EUCOMM via the European Mutant Mouse Archive (EMMA). Founder pups were generated in the UCSF transgenesis core facility by transgenic embryo implantation into pseudopregnant females. Pups were screened for WT *Il31* and mutant *Il31^tm1a(EUCOMM)Wtsi^* transgene alleles with the following PCR primer pairs:

*IL31 ^tm1a(EUCOMM)Wtsi^* CGAGAAGTTCCTATTCCGAAGT and GGAGGTTATCTGTGTCTCCATC

*IL31^WT^* allele GCAGCCAAAGATGTCCTAGT and TAGCCATCTGCAACCTGTTC

*IL31^null^* CGAGAAGTTCCTATTCCGAAGT and ACTTACTCAAGGTTATGCAGCT

### Other transgenic animals

All other mutant (knockout and transgenic) mouse strains used in the proposed research have been previously published and validated. These strains were acquired directly from the originating investigators (*Il31RA*^-/-^ from Genentech) or from Jackson Laboratories, as follows:

| Strain name | Official Symbol | Source | MGI ID | Pubmed |
| --- | --- | --- | --- | --- |
| IL31RA<sup>-/-</sup> | Il31ra<sup>tm1Ngh</sup> | Genentech | 3717594 | 17353366 |
| Flp/o | Gt(ROSA)26Sor | JAX 012930 | 104735 |  |
| β-actin-Cre | Tmem163<sup>Tg(ACTB-cre)2Mrt/EmsJ</sup> | JAX 033984 | 2176050 |  |

### Generation *of* Th1 and Th2 *in vitro-*differentiated CD4 T cell cultures

Bulk Th1-and Th2-polarized CD4 T cell cultures were generated by *in vitro-*differentiation of purified IL31KO or WT lymph node T cells, as previously described(*10, 48*). In brief, CD4 cells were isolated using the Easy-Sep CD4+ T cell isolation kit (Stemcell Technologies), plated on a tissue culture-treated plate pre-coated with neutravidin (Thermo), then stimulated for 3 days in DMEM media supplemented with 10% FBS (Omega Scientific), 1% L-glutamine, sodium pyruvate, and non-essential amino acids (Sigma), plus anti-mouse CD3 (clone 145-2C11; BioXCell) and anti-mouse CD28 (clone 37.51; BioXCell) which had been biotinylated using the EZ-link Sulfo-NHS-biotinylation kit (Thermo). Additionally, Th1 and Th2-polarizing cytokines and blocking antibodies were added as follows: For Th1, recombinant mouse IL-12 (Peprotech) and anti-IL-4 (BioXCell); for Th2, anti-IFNγ (XMG1.2, BioXCell) and IL-4 (10,000 U/ml stock added as a supernatant from I3L6 cells(*49*)).

### Chronic topical HDM sensitization

HDM ointment (Biostir AD, active ingredient *Dermatophagoides farinae*) for topical sensitization was obtained from Biostir, Inc. (Osaka, Japan). Sensitization was performed per published protocol(*17, 28*). In brief, shaved skin was treated twice-weekly with a 4% SDS in PBS solution followed 2h later by Biostir ointment application. For behavior and serum cytokine experiments cheek skin was sensitized for 8 weeks, and analysis performed in week 9. For flow cytometry and histology, dorsal neck skin was sensitized for 4 weeks then harvested in week 5. Toenails were clipped weekly from week 3 onward to prevent scratching-induced skin injury.

### OVA sensitization and intradermal challenge

Mice were sensitized on day 0 by intraperitoneal injection of 50 μg lyophilized BSA-free ovalbumin protein extract (Sigma) resuspended in PBS and pre-incubated with the adjuvant Imject Alum (Thermo). Mice received unilateral intradermal ear skin injections of 50 μg ovalbumin in 10 μl PBS daily on days 7-9.

### Skin histology

Paraffin embedding, tissue sectioning and H&E staining of skin was performed by the UCSF Mouse Pathology core. Images were captured with 10x and 20x objectives on a Zeiss AxioImager M2 microscope.

### RNA preparation and Taqman qPCR

For whole-skin RNA expression, RNA was extracted from skin using the RNeasy Fibrous Tissue kit (Qiagen). For bulk CD4 T cell RNA extraction, 10^6^ cells per sample were processed following the miRNeasy micro kit (Qiagen). After RNA quantification, cDNA was synthesized per the Invitrogen Superscript III kit protocol (Thermo). *Il31, Il4, Il13 and Ifng* cDNA abundance was quantified by qPCR using published Taqman primer/probe sets(*1, 50*).

### Skin digestion

Dorsal neck skin tissue sections were minced then digested for 1.5h in RPMI media (Sigma) supplemented with type XI collagenase (Sigma) and bovine deoxyribouclease I (Sigma). Enzymatic activity was quenched with fetal bovine serum (Omega Scientific) prior to mechanical dissociation by GentleMACS (Miltenyi). Resultant cell suspensions were filtered through 70 μm nylon fabric mesh.

### *Ex vivo* activation of skin T cells

When indicated, cells were activated by resuspension in DMEM media (Sigma) containing FBS (Omega Scientific), L-glutamine and Sodium Pyruvate (Sigma) and cultured for 4 hours at 37°C, 10% CO2 in the presence of phorbol 12-myristate 13-acetate, ionomycin, and Brefeldin A (Sigma).

### Flow cytometry

Cells were stained with DAPI or fixable viability dye eFluor-780 (eBioscience) and fluorophore-conjugated anti-mouse antibodies to cell surface antigens. For intracellular cytokine staining, cells were fixed in 4% paraformaldehyde (Electron Microscopy Sciences), then permeabilized in buffer from the eBioscience transcription factor staining kit (Thermo). Samples were collected on LSRII or LSRII Fortessa (Beckton-Dickinson) cytometers equipped with FACS Diva software. Flow cytometry data analysis was performed with Flowjo 10 software (Treestar).

Antibodies for flow cytometry included: BV711-conjugated anti-mouse CD45 (BD, clone 30-F11); AlexaFluor488-conjugated anti-mouse CD11b (eBiosciences; M1/70); PE-Cf594-conjugated anti-mouse CD3*ζ* (Biolegend, 145-2C11); PE-Cy7-conjugated anti-mouse CD4 (Biolegend, RM4-5); APC-conjugated anti-mouse CD4 (BioLegend; N418); BV605-conjugated anti-mouse CD8*α* (Biolegend, 53-6.7); BV605-conjugated anti-mouse *γδ*TCR (Biolegend, GL3) V450-conjugated anti-mouse Ly6C/G (BD Pharmingen; Gr-1); PE-conjugated anti-mouse SiglecF (BD Biosciences; E50-2440); APC-conjugated anti-mouse IL-4 (eBiosciences; 11B11) PE-e710-conjugated anti-mouse IL-13 (eBiosciences; eBio13A); eFluor450-conjugated anti-mouse IFNγ (eBiosciences; XMG1.2); Alexa Fluor 488-conjugated anti-mouse IL-17A (eBiosciences; eBio17B7); PE-conjugated anti-mouse IL-31 (Biolegend; W17037B).

### RNA preparation for scRNA-seq

HDM-treated dorsal neck skin from 4 pooled Il31RAKO and 4 pooled WT C57Bl/6 mice was digested and prepared for flow cytometry as above. The live (DAPI-) CD45+ cell fraction of each sample was sorted using a Moflo XDP (Beckman Coulter) directly into ice-cold 0.5% BSA in PBS, and immediately processed through the Chromium Single Cell 3′ v2 Library Kit (10X Genomics) per the manufacturer’s protocol by the UCSF Genomics Core Facility of the UCSF Institute for Human Genetics. Each channel was loaded with 30,000 cells per sample. The cells were then partitioned into Gel Beads in Emulsion in the instrument, where cell lysis and barcoded reverse transcription of RNA occurred, followed by amplification, shearing, and 5′ adaptor and sample index attachment. Libraries were sequenced on an Illumina HiSeq 4000. Single Cell 3′ libraries used standard Illumina sequencing primers for both sequencing and index reads and were run using paired-end sequencing with single indexing where Read1 is 26 cycles and Read 2 is 98 cycles.

### scRNA-seq Analysis

Initial post-processing and quality control including single-cell transcript alignment to mouse genome GRC38/mm10 were also performed by the UCSF Genomics Core Facility, using the 10X Cell Ranger package (Cell Ranger Version 3.1, 10X Genomics).

R package Seurat (version 3.2.2, Satija Lab) was used for single cell transcriptome analysis. Expression matrices for WT and IL31RA were combine using the Seurat merge command. The SCTransform normalization method was used to control for between-sample sequencing depth variation. Cells were removed according to the following thresholds: <500 genes/cell or >5000 genes per cell, >20000 UMIs/cell, 10% mitochondrial content, <0.01% hemoglobin content. Among the cells retained, the effects of mitochondrial and ribosomal content were regressed out prior to clustering. Clustering was performed using the default Seurat parameters: clusters are identified using Shared Nearest Neighbors method then optimized using the original Louvain algorithm. Genes were excluded from the final dataset if they were expressed in fewer than 3 cells, resulting in a total of 15,592 genes. For differential expression (DEG) analysis between clusters, genes were detected if they are expressed in at least 10% of cells in a cluster with a log fold change of at least 0.25. For DEG analysis between comparison groups, genes were detected if expressed in at least 5% of cells in the group with a log fold change of at least 0.25.

Myeloid cell clusters identified by inspection of the initial Seurat DEGs were selected to generate a Myeloid Seurat object using Seurat’s subset command, and re-clustering was performed using SCTransform parameters as above. These DEGs were used as cell marker genes to assign myeloid cluster identities (Fig. S5). Neutrophils and a cluster containing predominantly T cell-myeloid doublets (identified by co-expression of genes encoding subunits and Cd209/DC-SIGN) were removed from subsequent analyses. For comparative analysis within macrophages or dendritic cells, as depicted in Fig. 4D-E, additional Seurat objects were generated from the Myeloid Seurat object, and cluster markers genes for these new Seurat objects were identified using the same computational workflow and DEG detection parameters described above.

### IgE and IL-4 ELISAs

For serum IgE quantification, MaxiSorp 96-well plates (Nunc) were prepared by coating with anti-mouse IgE antibody (R35-72, BD) then blocking with 1% BSA in PBS. Serum samples (1:25 dilution) and a serially-titrated mouse IgE standard (C48-2, BD) were added to the plate and incubated for 2h. Plates were washed with PBST, incubated with biotin-conjugated anti-mouse IgE secondary antibody (R35-118, BD), washed again, and incubated with HRP Streptavidin (Biolegend). For serum IL-4 quantification, plates were coated with anti-mouse IL-4 antibody (11B11, Biolegend) then blocked as above. Serum samples (1:4 dilution) and a serially-titrated mouse IL-4 standard (Peprotech) were added and incubated for 2h. Plates were washed, incubated with biotin-conjugated anti-mouse IL-4 secondary antibody (BVD6-24G2, BD), washed again, and incubated with HRP Streptavidin (Biolegend). After final washes, plates were developed with TMB substrate (Invitrogen), stopped with 2N H2SO4, and read at 450nm on a microplate reader (Molecular Devices).

### Behavior monitoring and analysis

Scratching bouts per 30-minute window of videorecording were tabulated by a blinded observer, following a standard protocol validated by Lamotte(*51*). For HDM-treated animals, spontaneous scratching bouts were tabulated in week 5 and week 9, and at least 2 days following the previous HDM treatment, as previously described(*28*). For acute scratching, nape-of-the-neck intradermal injections of 100 ul PBS or solutions containing synthetic pruritogens were performed immediately prior to videorecording. Injected pruritogens included histamine (500 μg; Sigma), chloroquine (200 μg; Sigma), SLIGRL (100 nmol; Peptides International), *α*-methyl-5-hydroxytryptamine (500 μg; Abcam), and endothelin-1 (10 pmol; Calbiochem).

### RNAscope *in situ* hybridization

*In situ* hybridization was performed on flash-frozen sections of trigeminal ganglia using the RNAscope Fluorescent Multiplex in situ hybridization kit (ACD Biosciences), per manufacturer’s instructions. RNAscope probes (ACD Biosciences) included: IL31RA (C1, C2 and C3); OMSR (C3); TRPV1 (C1, C2 and C3); TRPA1 (C2); IL4RA (C1 and C2); F2RL1 (C1, C2 and C3), with Alexa-488, Atto-550 and Atto-647 dyes.

### *Ex vivo* calcium imaging of DRG neurons

Lumbar spinal cord DRG from 6–10 week-old WT and IL31KO mice were collected, enzymatically-digested with collagenase II (Worthington), dispase II (Sigma) and papain (Worthington), and mechanically triturated as previously described(*52*). The resultant single cell suspension was plated on poly-D-lysine/laminin-coated coverslips (Corning Biocoat), incubated at 37 °C in 95% humidity (95%) and 5% CO_2_, and imaged within 16–36 h.

DRG cells were loaded with Fura-2-AM and 0.01% Pluronic F-127 (Invitrogen), then mounted in the open chamber of a Nikon Eclipse TE300 microscope for imaging. Cells were sequentially challenged by addition of the following agonist solutions: SLIGRL (100 μM; R&D Systems), then either rmIL-31 (300 nM; Peprotech) or BAM8-22 (2 μM; Tocris), then capsaicin (1 μM; Sigma), followed by KCl (50 mM; Sigma) to distinguish neurons from non-neural cells.

Fura-2-AM fluorescence was measured at 340 nm and 380 nm excitation and 530 nm emission. COOL LED PE300 was used as the excitation source. Images were acquired using a Nikon TE300 microscope fitted with a Hamamatsu ORCA-ER camera and MetaFluor v7.6.9 software (Molecular Devices). Images were analyzed using a custom journal in MetaMorph v7.6.9 software (Molecular Devices). Increased 340/380 ratio ≥0.2 over baseline counted as a positive response to agonist.

### Institutional ethics approval

Euthanasia procedures and experimental protocols were approved by the University of California, San Francisco IACUC committee (protocols AN183584-01B and AN183265-01E), and performed in accordance with University of California, San Francisco IACUC Committee guidelines and regulations.

### Statistical testing and analysis

Data were analyzed using Prism 8 software (Graphpad) by comparison of means testing using unpaired two-tailed student’s t-tests unless otherwise indicated. Error bars indicate mean ± standard deviation. All data points reflect unique biological replicates; no technical replicates or repeated measurements were performed. For experiments where data collection required manual counting (behavior analysis, cell counting in *in situ* hybridization slides), scoring was performed by a lab member blinded to sample identity, genotype, and intervention/control.

### Cohort composition and sample sizes

WT and KO animals in our colony were age-and sex-matched, then randomly assigned to control vs treatment groups. For all experiments, we used mixed cohorts of male and female mice (aged 8-12 weeks for in vivo experiments; aged 4.5-6 weeks for *in vitro* T cell polarization assays). In age-matching, a maximum tolerance of 2 weeks age difference was considered acceptable. A power calculation was used to determine sample sizes needed for in vivo dermatitis models. Groups of 5-6 mice/HDM or OVA treatment group were used, providing <10% chance of improper rejection of the null hypothesis (no difference in the mean frequency of infiltrating inflammatory cells in any two sets of mice), assuming the difference to be measured is (at minimum) a doubling of the number of rare inflammatory immune cell subsets from 1% to 2% with a standard error of 0.5% in each group. These assumptions are consistent with our own observations and with published data on dermatitis models.

## Supplementary Materials

Figure S1. Transgene schematic and serial crosses to generate *Il31^null^* allele

Figure S2. Supplementary data to accompany Figure 2

Figure S3. Gating strategy for cutaneous CD45+ cell subsets

Figure S4. Gating strategy for CD4 T cell intracellular cytokine detection

Figure S5. Cell marker genes for the 13 myeloid clusters

Figure S6. Supplementary data to accompany Figures 4-5

## Acknowledgments

We acknowledge Nico Ghilardi of Genentech (IL31RAKO mice), Biolegend (early access to anti-mouse IL-31 antibody), and additional members of the Ansel and Basbaum laboratories.

## Funding

Dermatology Foundation Career Development Award (MSF)

National Institute of Arthritis and Musculoskeletal and Skin Diseases K08AR074556 (MSF) National Heart, Lung, and Blood Institute R01HL109102 (KMA)

Sandler Foundation (KMA)

National Institute of Neurological Disorders and Stroke R35NS097306 (AIB) National Institutes of Health P30 DK63720 (UCSF Parnassus Flow Cytometry Core)

## Author contributions

Conceptualization: MSF, KMA, AIB

Methodology: MSF, JMB, CC, KMA, AIB

Investigation: MSF, JMB, CC, MS, DM, CC

Visualization: MSF, JMB, AWS

Funding acquisition: KMA, AIB

Supervision: KMA, AIB

Writing – original draft: MSF, KMA

Writing – review & editing: JMB, CC, JS, AIB, KMA

## Data and materials availability

All data described in this manuscript will be made available upon request to the corresponding author. Prior to publication, raw sequencing files will be deposited in GEO at NCBI. ES cells containing the *Il31* targeted allele are available from EUCOMM via the European Mutant Mouse Archive. IL31RAKO mice were obtained by MTA from Genentech and cannot be distributed by the authors. All other materials are available directly from commercial suppliers.

