## Supplemental Figures for "IL-31 uncouples skin inflammation from itch sensation in allergic dermatitis"

**Supplementary Material:**

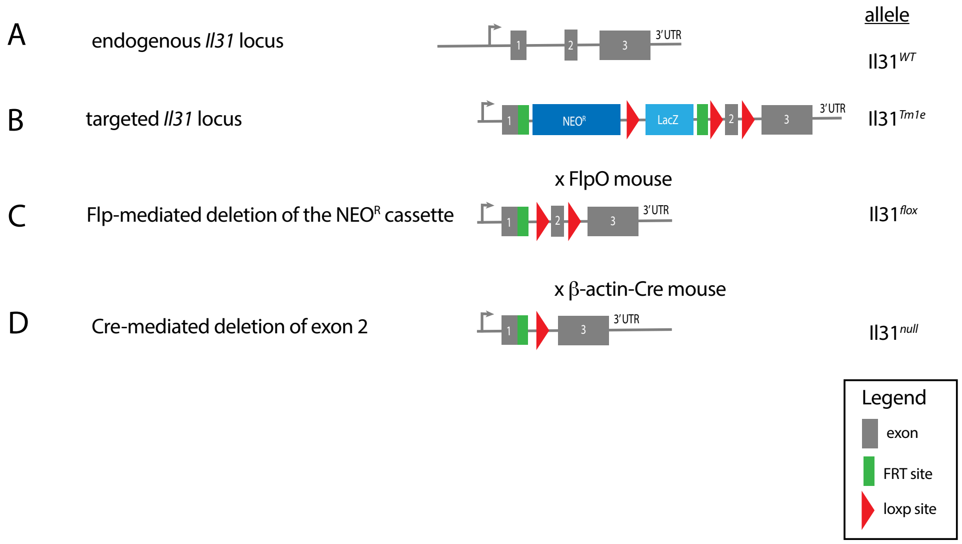


**Fig. S1**. **Transgene schematic and serial crosses to generate** ***Il31*****null allele**

**(A)** Endogenous 3-exon *Il31* allele

**(B)** The *Il31Tm1e* transgene interrupts exon 1 of the endogenous *Il31* locus with a neomycin resistance/LacZ cassette.

**(C)** Mice containing an *Il31flox* allele were generated by crossing an *Il31Tm1e* transgenic mouse with a Flp-recombinase expressing strain (FlpO), resulting in germline excision of the region between FRT sites (green rectangles).

**(D)** Mice containing an *Il31null* allele were generated by crossing an *Il31flox* transgenic mouse with a ubiquitous Cre recombinase-expressing strain (β-actin Cre), resulting in germline excision of the region between loxp sites (red triangles).

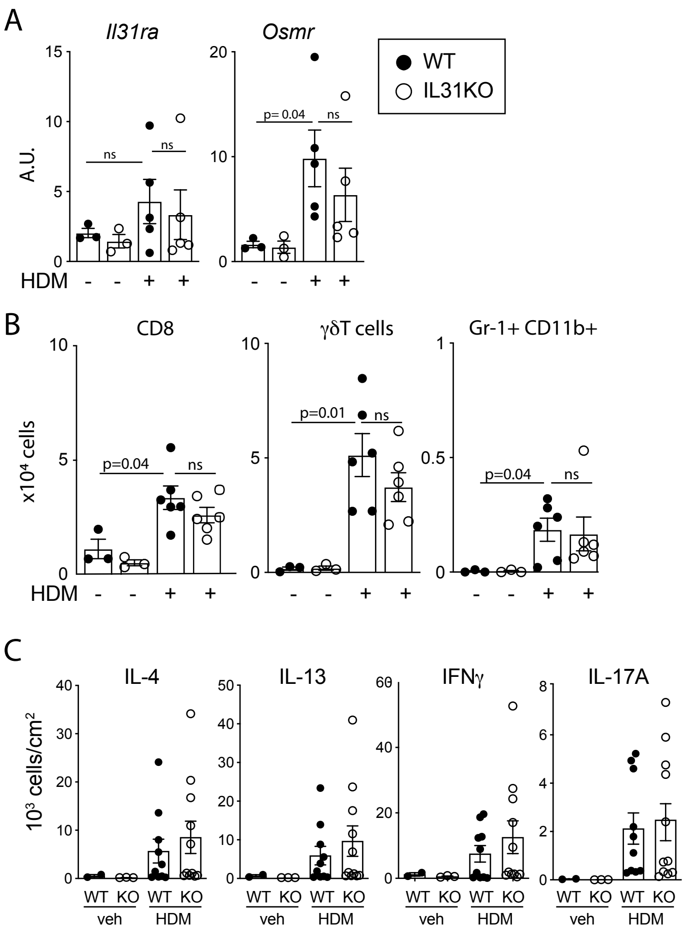


**Fig. S2**. **Supplementary data to accompany Figure 2**

**(A)** Taqman-based qPCR detection of *Il31ra* and *Osmr* in control and HDM-treated skin.

**(B)** CD45+ (hematopoietic) populations increased in HDM-treated skin include CD8 T cells, dermal γδ T cells, and CD11b+Gr1+ cells (combined monocytes and neutrophils).

**(C)** Flow cytometry data from Figure 2F enumerated as cytokine-producing CD4 T cells per mouse. Both vehicle (control) and HDM-treated samples are included for comparison.

WT, black circles; IL31KO, open circles. Data reflect n=3 animals per control group and n≥5 per HDM treatment group, and are representative of 3 independent experiments (A, B), or pooled from 2 independent experiments with n≥5 mice per treatment group (C). Bars indicate mean +/- SD. Statistical significance determined by unpaired student's t-test.

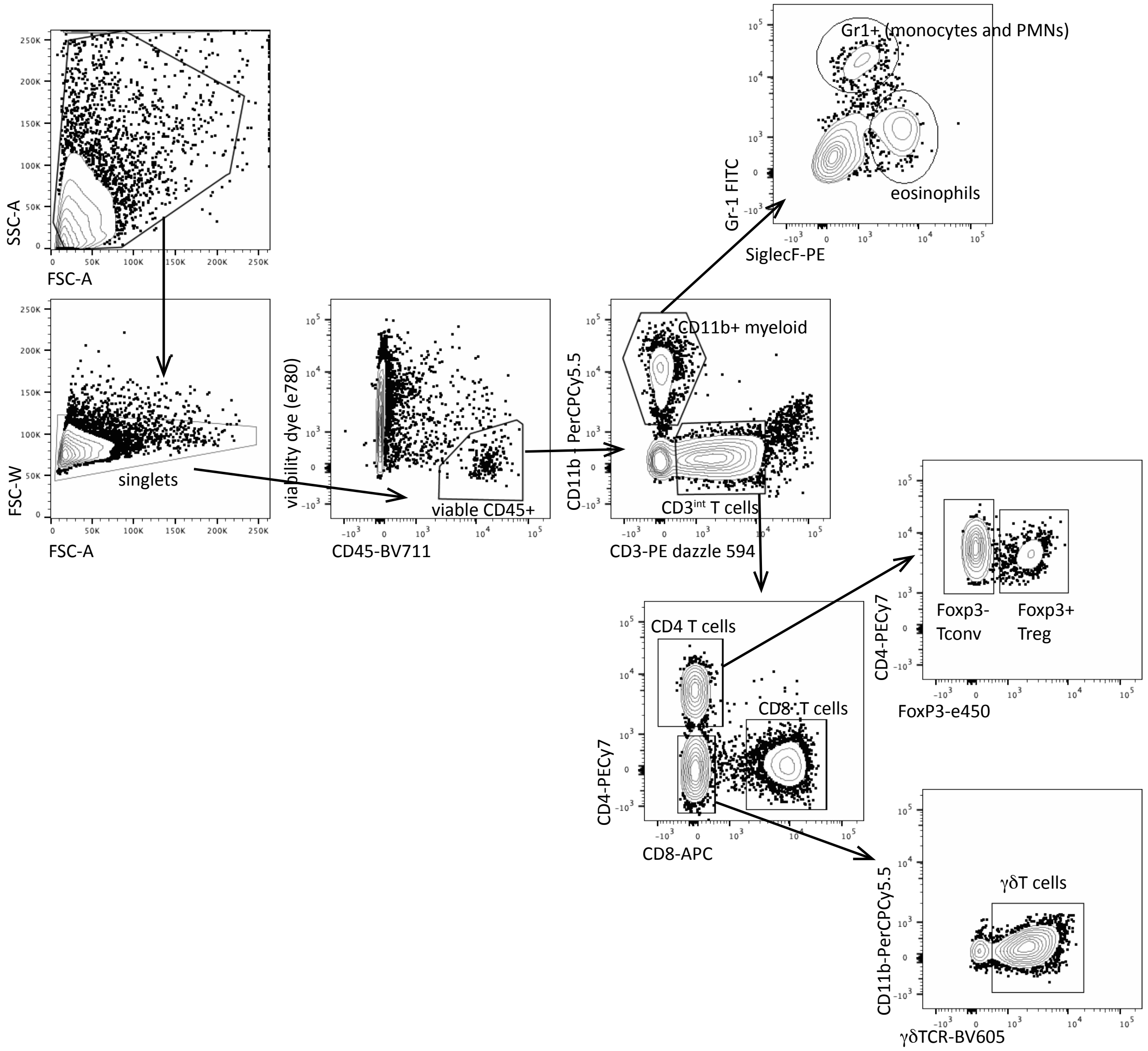


**Fig. S3**. **Gating strategy for cutaneous CD45+ (hematopoietic) subsets**

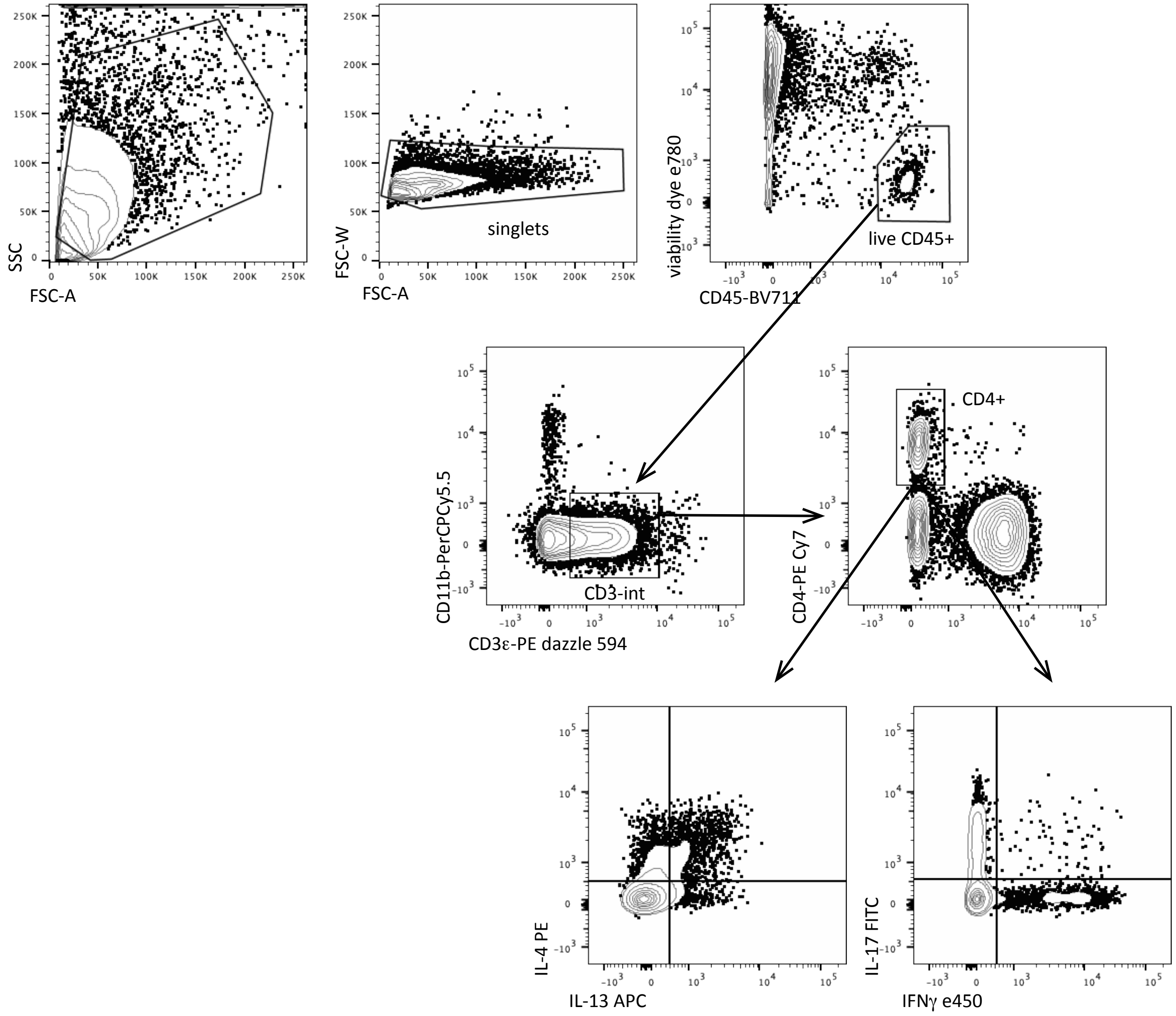


**Fig. S4**. **Gating strategy for CD4 T cell intracellular cytokine detection**

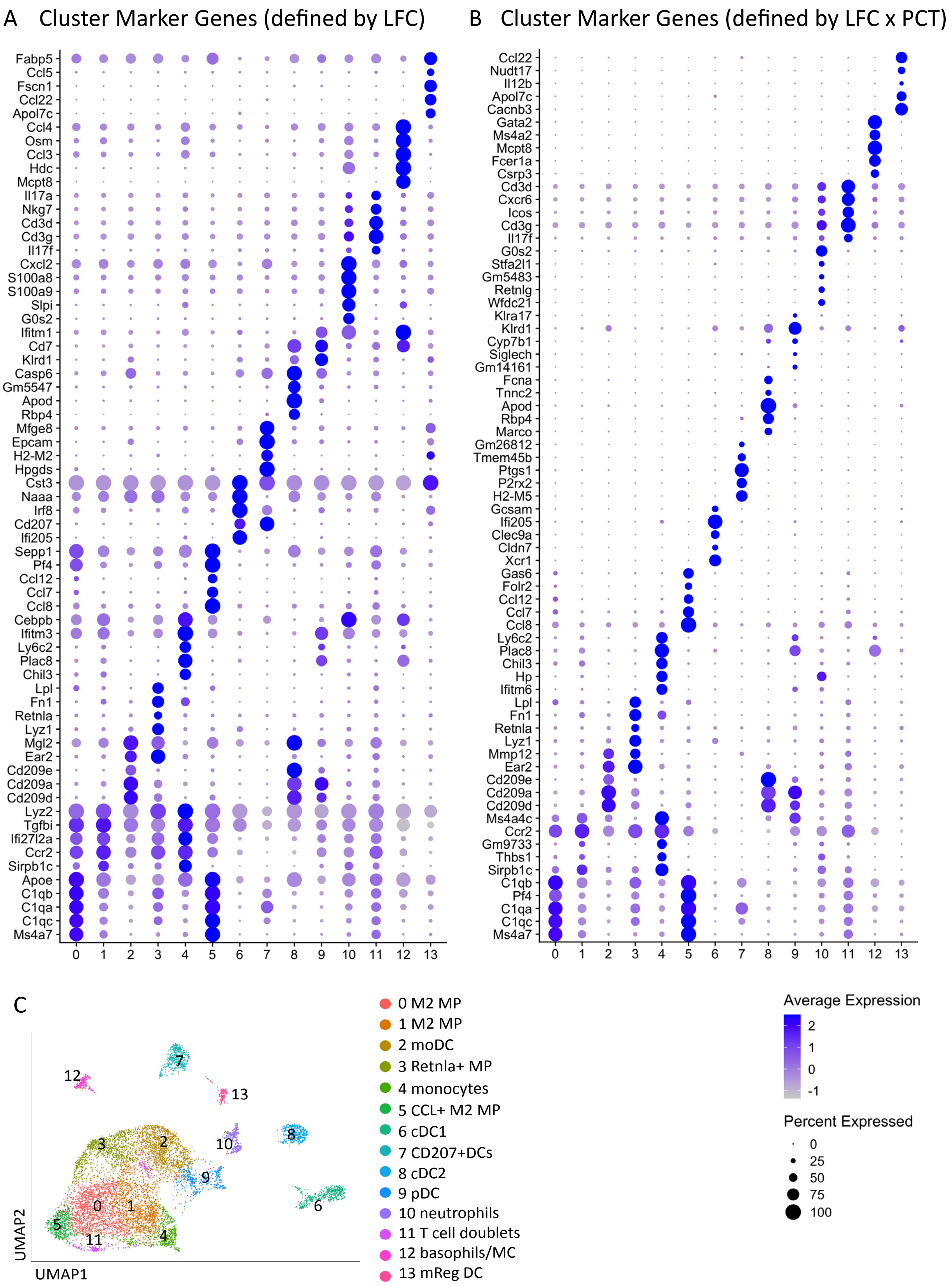
A Cluster Marker Genes (defined by LFC)
Fabp5
Ccl5
Fscn1
Ccl22
Apol7c
Ccl4
Osm
Ccl3
Hdc
Mcpt8
Il17a
Nkg7
Cd3d
Cd3g
Il17f
Cxcl2
S100a8
S100a9
Slpi
G0s2
Ifitm1
Cd7
Klrd1
Casp6
Gm5547
Apod
Rbp4
Mfge8
Epcam
H2-M2
Hpgds
Cst3
Naaa
Irf8
Cd207
Ifi205
Sepp1
Pf4
Ccl12
Ccl7
Ccl8
Cebpb
Ifitm3
Ly6c2
Plac8
Chil3
Lpl
Fn1
Retnla
Lyz1
Mgl2
Ear2
Cd209e
Cd209a
Cd209d
Lyz2
Tgfbi
Ifi27l2a
Ccr2
Sirpb1c
Apoe
C1qb
C1qa
C1qc
Ms4a7
0 1 2 3 4 5 6 7 8 9 10 11 12 13
B Cluster Marker Genes (defined by LFC x PCT)
Ccl22
Nudt17
Il12b
Apol7c
Cacnb3
Gata2
Ms4a2
Mcpt8
Fcer1a
Csrp3
Cd3d
Cxcr6
Icos
Cd3g
Il17f
G0s2
Stfa2l1
Gm5483
Retnlg
Wfdc21
Klra17
Klrd1
Cyp7b1
Siglech
Gm14161
Fcna
Tnnc2
Apod
Rbp4
Marco
Gm26812
Tmem45b
Ptgs1
P2rx2
H2-M5
Gcsam
Ifi205
Clec9a
Cldn7
Xcr1
Gas6
Folr2
Ccl12
Ccl7
Ccl8
Ly6c2
Plac8
Chil3
Hp
Ifitm6
Lpl
Fn1
Retnla
Lyz1
Mmp12
Ear2
Cd209e
Cd209a
Cd209d
Ms4a4c
Ccr2
Gm9733
Thbs1
Sirpb1c
C1qb
Pf4
C1qa
C1qc
Ms4a7
0 1 2 3 4 5 6 7 8 9 10 11 12 13
C
UMAP2
UMAP1
0 M2 MP
1 M2 MP
2 moDC
3 Retnla+ MP
4 monocytes
5 CCL+ M2 MP
6 cDC1
7 CD207+DCs
8 cDC2
9 pDC
10 neutrophils
11 T cell doublets
12 basophils/MC
13 mReg DC
Average Expression
2
1
0
-1
Percent Expressed
0
25
50
75
100

**Fig. S5**. **Cell Marker genes for the 13 myeloid clusters**

**(A)** Dot plot of top 5 differentially-expressed genes defined by average log fold change (LFC)

**(B)** Dot plot of top 5 differentially-expressed genes defined as the product of the average log fold change (LFC) and the cluster specificity (PCT ratio).

**(C)** UMAP plot and cluster identities from Figure 4 reproduced for reference

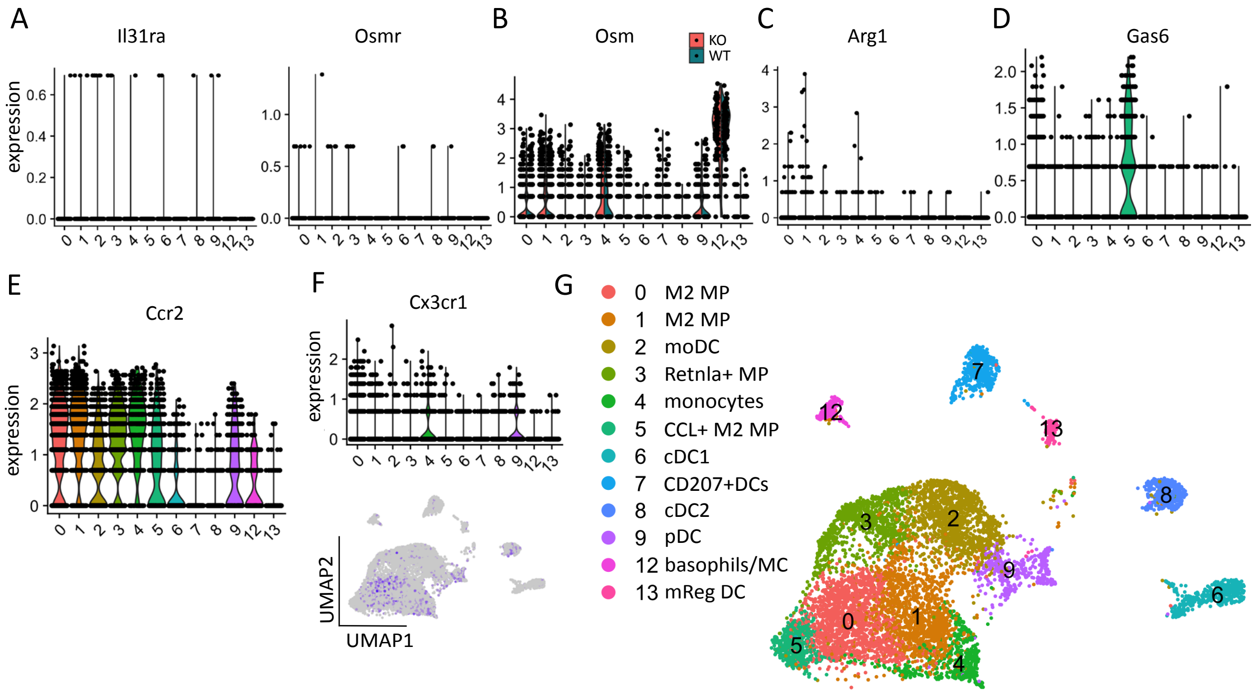


**Fig. S6. Supplementary data to accompany Figures 4-5.**

**(A)** *Il31ra* and *Osmr* only detected in scattered cells

**(B)** Split violin plot demonstrates similar *Osm* expression in WT and IL31RAKO across clusters (turquoise = WT, salmon = IL31RAKO).

**(C)** Rare cells express M2 marker *Arg1*

**(D)** Expression of Mertk ligand and cluster 5 marker *Gas6*.

**(E)** *Ccr2* is broadly expressed by macrophages (clusters 0,1,3,4,5) and monocyte-derived DCs (cluster 2).

**(F)** Distribution of *Cx3cr1* across myeloid clusters (top: violin plot; bottom, feature plot)

**(G)** UMAP plot and cluster identities from Figure 5 reproduced for reference
